# Quantitative Profiling Method for Oxylipins in Neurodegenerative Diseases by Liquid Chromatography Coupled with Tandem Mass Spectrometry

**DOI:** 10.1101/2023.10.02.560544

**Authors:** Elham Pourmand, Fan Zhang, Morteza Sarparast, Jamie K. Alan, Kin Sing Stephen Lee

**Affiliations:** Department of Chemistry, Michigan State University, East Lansing, MI, USA; Department of Pharmacology and Toxicology, Michigan State University, East Lansing, MI, USA; Institute of Integrative Toxicology, Michigan State University, East Lansing, MI, USA

## Abstract

Aging is one of the major risk factors for many chronic diseases, including diabetes, neuropathy, hypertension, cancer, and neurodegenerative diseases. However, the mechanism behind aging and how aging affects a variety of disease progression remains unknown. Recent research demonstrated the cytochrome P450 (CYP)-epoxide hydrolase (EH) metabolites of polyunsaturated fatty acids (PUFAs) play a critical role in the abovementioned age-associated diseases. Therefore, aging could affect the abovementioned chronic diseases by modulating CYP-EH PUFA metabolism. Unfortunately, investigating how aging affects CYP-EH metabolism in human and mammalian models poses significant challenges.

In this regard, we will use *C. elegans* as a model organism to investigate the aging effects on CYP-EH metabolism of PUFA, owing to its long history of being used to study aging and its associated benefits of conducting aging research. This project will develop analytical tools to measure the endogenous levels of CYP-EH PUFA metabolites in *C. elegans* using state-of-the-art ultra-performance liquid chromatography coupled with tandem mass spectrometry (UPLC-MS/MS). These metabolites are very potent but present in low abundance. The dramatic increase in sensitivity in UPLC-MS/MS allows us to monitor these metabolites over the lifespan of C. *elegans* with minimum samples. Our results show that *C. elegans* produces similar CYP PUFA metabolites to mammals and humans using our SPE-UPLC-MS/MS method. We will also show that our method successfully determined the CYP-EH PUFA metabolites profile changes induced by the inhibition of *C. elegans* EH. The method developed from this project will significantly improve our understanding of the role of dietary PUFAs and associated metabolism on aging and neurodegeneration and will uncover new mechanisms of how aging affects neurodegeneration through the modulation of PUFA metabolic pathways.

**Graphical abstract:** 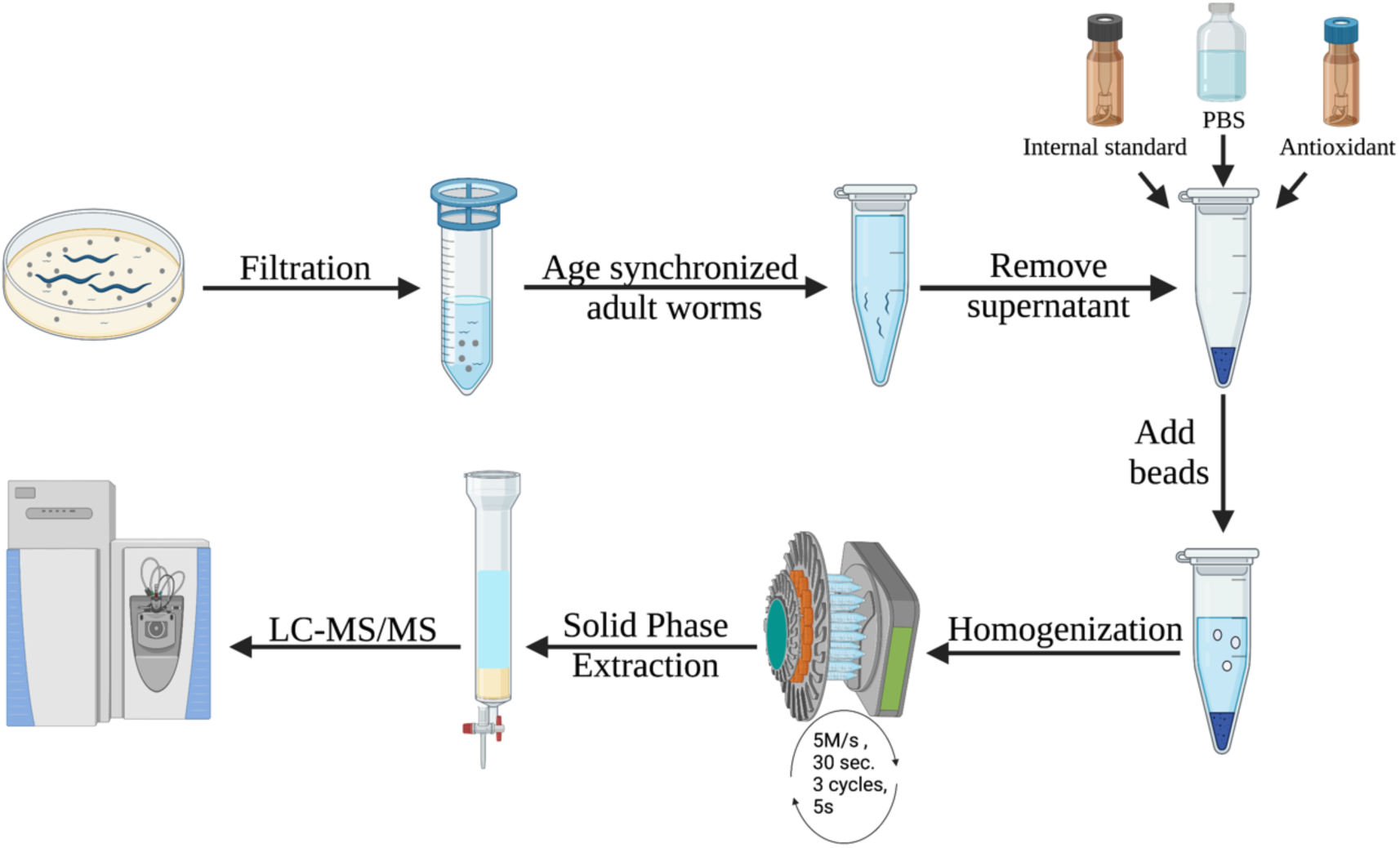

## 1. Introduction

Aging is one of the major risk factors for many diseases, particularly neurodegenerative diseases. For example, it is projected that by 2050, the prevalence of Alzheimer’s Diseases (AD) will double every 5 years after age 65, ^1^ and the number of patients with AD will triple by 2050 without an effective treatment. ^2^ Thus, understanding and modifying the aging process could hugely impact patients with AD and other aging-associated neurodegenerative diseases. Recent research showed that dietary lipids have significant effects on aging and cell senescence, ^3–5^ and their downstream oxidized polyunsaturated fatty acid (PUFA) metabolites (namely oxylipins) are key lipid mediators for many disease states including but not limited to, cancer, lupus, cardiovascular^6-8^, and neurodegenerative diseases.^9-12^An increase in the expression of enzymes that produce oxylipins, including, lipoxygenase (LOX) ^13^, Cyclooxygenase (COX)^14^ soluble epoxide hydrolase (sEH), and microsomal epoxide hydrolase (mEH) is observed in AD compared to healthy individuals. ^15-17^ As a result, endogenous levels of specific oxylipins produced by sEH are upregulated in AD individuals. Moreover, inhibiting the epoxide hydrolase (EH) metabolism of epoxy fatty acids, to their dihydroxy fatty acid metabolites by inhibitors or a genetic knockout is beneficial for neurodegenerative diseases, including Parkinson’s disease (PD) and AD. ^18,19^ (**Figure 1**) Our recent study also indicated that dihydroxyeicosadienoic acid (DHED), an epoxide hydrolase metabolite of dihomo-gamma-linolenic acid, induces degeneration of dopaminergic neurons through ferroptosis, which could affect aging.^11^ However, the exact role of oxylipins in neurodegeneration and whether they play any critical role in aging remains elusive. ^20^ Therefore, investigating whether aging has any effects on the oxylipins level and *vice versa* will help to develop more effective preventative measures and treatments for age-related diseases, including neurodegenerative diseases. However, investigating the molecular interactions between oxylipins and aging poses several challenges. First, the mechanisms of aging are understudied and complex. Therefore, conducting studies on an intact animal will likely be beneficial. In addition, aging is a long process, and the experimental settings are difficult to control throughout the aging studies. ^21-23^ Moreover, oxylipins are largely present at low levels, and there are hundreds of different oxylipins produced in animals. To further complicate the picture, the structures of these oxylipins are very similar, but their functions are very diverse and sometimes in opposition. ^24^ Besides, the endogenous levels for some oxylipins can change up to 3 orders of magnitude depending on the disease status. ^25^ Therefore, it is necessary to monitor the *in vivo* level of every oxylipin to understand how oxylipin biosynthesis affects the aging process comprehensively. Yet, this creates very difficult analytical challenges.

**Figure 1.**
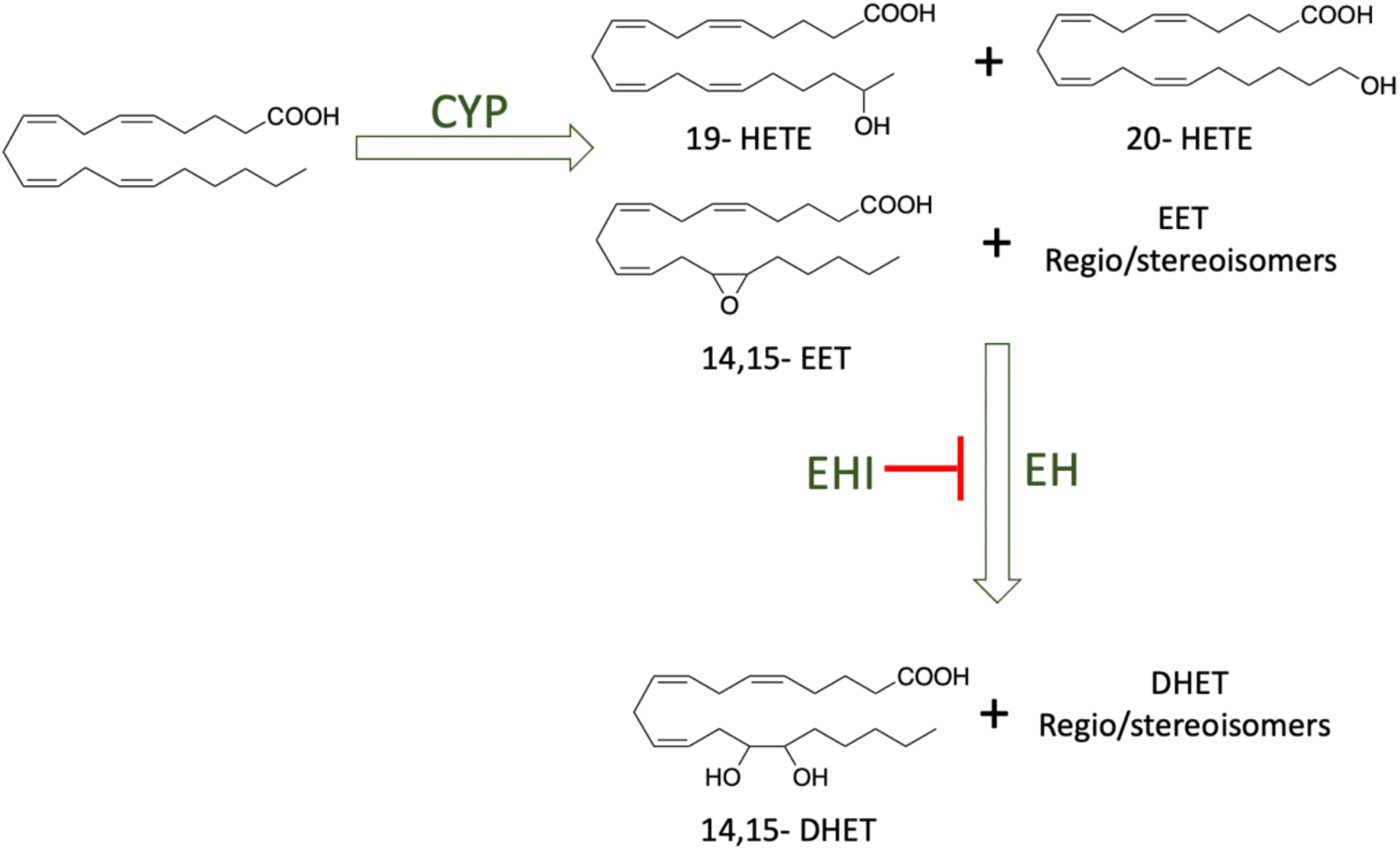
Metabolic pathway of polyunsaturated fatty acids. Cytochrome 450 (CYP), epoxide hydrolase (EH), and epoxide hydrolase inhibitor (EHI).

To solve these challenges, this study used *Caenorhabditis elegans* (*C. elegans*) as an animal model to investigate the molecular interaction between aging and oxylipin biosynthesis. *C. elegans* has been used for aging research for decades, owing to its short lifespan, ease of handling and ability to conduct imaging studies, adaptability to high-throughput assays, possessing similar neuronal structures and functions, and the presence of a conserved genome between *C. elegans* and humans. ^26^ In addition, we developed an analytical method using ultra-performance liquid chromatography (HPLC) coupled with tandem mass spectrometry to monitor and quantify the oxylipin profile in *C. elegans.* The developed method is a combination of different analytical techniques, including solid phase extraction (SPE), HPLC, electrospray ionization (ESI), and tandem mass spectrometry to tackle all the challenges associated with qualification and quantification of oxylipins in *C. elegans*. Our results indicate that we successfully developed a fully validated analytical method to quantify the oxylipins in *C. elegans*. Furthermore, we determined that *C. elegans has* a conserved cytochrome P450-epoxide hydrolase metabolic pathway for PUFAs. Our method will eventually facilitate an understanding of the biological functions of oxylipins in aging.

## 2. Results and discussion

### 2.1. Mass spectrometry optimization

Optimization of the LC and MS parameters was done to improve oxylipin detection because these molecules are present in very low concentrations. Considering this, the mass parameters, including collision and cone voltage, were optimized to increase the signal-to-noise ratio, while simultaneously decreasing the limit of detection (LOD). In a quadrupole-based mass spectrometer, dwell time, which is the amount of time a mass spectrometer spends measuring the intensity of a particular mass-to-charge ratio during a single scan, controls the frequency of data acquisition. Thus, the dwell time could be considered as a trade-off between sensitivity and noise. While the longer dwell time provides a higher number of measurements across a peak, experiencing higher noise is inevitable. Thus, to get the maximum number of measurements with minimum noise, the acquisition was divided into ten functions based on the retention time of the compounds. We employed the “auto” feature available in the MassLynx V4.2 software, which automatically determined the dwell time by considering the number of transitions in each function. By utilizing this approach, we ensured sufficient time for acquiring reliable signal-to-noise ratios and an appropriate number of scans per peak. Furthermore, a type I internal standard (IS) was used to normalize the loss of oxylipins due to sample preparation. Type I internal standards are essentially deuterated versions of selected oxylipins that can be easily differentiated by mass spectrometry. The rationale for selecting such internal standards lies in their similarity to the physical properties of the target compounds. This similarity ensures that the loss experienced during extraction is consistent, allowing for the generalization of the loss to other oxylipins, therefore making an accurate normalization. Another advantage of using deuterated oxylipins as internal standards is that their endogenous concentrations in biological samples are negligible. As a result, the signal observed in mass spectrometry analysis is solely derived from the spiked internal standard with known concentration. A list of ISs, their concentration, and more detail is provided in **(SI Table 1).**

To achieve optimal sensitivity and selectivity in this method, a thorough analysis was conducted to identify specific fragments that occur in the ionized molecules of the sample. The selection of most transitions was based on identifying cleavages near double bonds or functional groups like epoxy or hydroxyl modification. In certain cases, prioritizing selectivity over sensitivity was deemed crucial, leading to the selection of a more distinctive fragmentation ion, even if it wasn’t the most intense. This approach ultimately contributes to a more precise analysis. For instance, in the case of the CYP450 metabolite of EPA, 8,9-EEQ, a fragment with a m/z of 127 was chosen instead of the more sensitive fragment at 255, with the primary aim of enhancing the selectivity of analysis of these regioisomers at the presence of 11,12-EEQ. Selecting m/z of 127 is crucial since these two isomers have very close retention times and the exact same molecular weight **(SI Figure 2).** The summary of optimized transitions and mass spectrometric parameters is listed in **Table 1**. By meticulously selecting suitable multiple reaction monitoring (MRM) transitions, optimizing cone and collision energy, and employing the appropriate LC mobile phases and gradient (**SI Table 3),** ^28^ we achieved successful separation and detection of distinct metabolites of PUFAs and regioisomers (**SI Figure 2),** With all these considerations, the total number of oxylipins in this method was higher compared to other studies. ^28,29^

**Table 1.** Summary of multiple reaction monitoring (MRM)(m/z), collision energy (CE), cone voltage (CV), retention time (RT)(min), the lower limit of quantification (LLOQ) (nM), the limit of detection (LOD)(nM), the upper limit of quantification (ULOQ)(nM) of oxylipin metabolite.

| Analytes | MRM | Cone Volt. | Collision Volt. | RT (min) | LLOQ (nM) | LOD (nM) | ULOQ (nM) |
| --- | --- | --- | --- | --- | --- | --- | --- |
| 15,16-DiHODE | 311.20 > 223.00 | 44 | 22 | 6.31 | 0.063 | 0.021 | 50 |
| 12,13-DiHODE | 311.20 > 183.00 | 36 | 22 | 6.33 | 1.250 | 0.416 | 1000 |
| 9,10-DiHODE | 311.20 > 201.00 | 52 | 16 | 6.35 | 0.063 | 0.021 | 25 |
| 17,18-DiHETE | 335.20 > 247.00 | 39 | 16 | 6.63 | 0.139 | 0.046 | 1000 |
| 14,15-DiHETE | 337.20 > 207.00 | 33 | 16 | 6.97 | 0.063 | 0.021 | 25 |
| 11,12-DiHETE | 335.10 > 167.00 | 36 | 16 | 7.15 | 0.063 | 0.021 | 25 |
| 12,13-DiHOME | 313.20 > 183.00 | 20 | 22 | 7.30 | 0.063 | 0.021 | 50 |
| 8,9-DiHETE | 335.20 > 127.00 | 36 | 20 | 7.50 | 0.250 | 0.083 | 100 |
| 9,10-DiHOME | 313.20 > 201.00 | 20 | 22 | 7.70 | 0.063 | 0.021 | 50 |
| 14,15-DiHETrE | 337.20 > 207.00 | 33 | 16 | 8.00 | 0.063 | 0.021 | 50 |
| 11,12-DiHETrE | 337.20 > 167.00 | 51 | 22 | 8.60 | 0.250 | 0.083 | 200 |
| 20-HEPE | 317.20 > 243.00 | 28 | 16 | 8.65 | 0.625 | 0.208 | 500 |
| 14,15 DHED | 339.10 > 209.00 | 28 | 22 | 8.84 | 0.125 | 0.042 | 100 |
| 9-HOTrE | 293.20 > 171.00 | 40 | 16 | 8.84 | 0.312 | 0.104 | 125 |
| 10,11-DiHDPE | 361.20 > 153.00 | 46 | 16 | 8.91 | 0.125 | 0.042 | 100 |
| 13-HOTrE | 293.20 > 195.00 | 36 | 16 | 9.02 | 0.625 | 0.208 | 250 |
| 18-HEPE | 317.10 > 215.00 | 44 | 16 | 9.03 | 0.625 | 0.208 | 500 |
| 8,9-DiHETrE | 337.20 > 127.00 | 15 | 22 | 9.18 | 0.250 | 0.083 | 200 |
| 11,12-DHED | 339.10 > 169.00 | 28 | 22 | 9.25 | 0.083 | 0.028 | 100 |
| 19-HETE | 319.10 > 275.00 | 44 | 16 | 9.44 | 0.625 | 0.208 | 250 |
| 5,6-DiHETE | 335.20 > 145.00 | 21 | 16 | 8.20 | 10.000 | 5.000 | 2000 |
| 15-HEPE | 317.00 > 219.00 | 36 | 10 | 9.52 | 0.312 | 0.035 | 250 |
| 20-HETE | 319.20 > 275.00 | 44 | 16 | 9.58 | 0.312 | 0.104 | 250 |
| 8,9-DHED | 339.10 > 187.00 | 28 | 22 | 9.63 | 0.125 | 0.042 | 100 |
| 7,8-DiHDPE | 361.20 > 113.00 | 44 | 22 | 9.64 | 0.625 | 0.208 | 250 |
| 8-HEPE | 317.10 > 155.00 | 36 | 16 | 9.91 | 0.208 | 0.069 | 100 |
| 12-HEPE | 317.10 > 179.00 | 44 | 10 | 9.91 | 0.312 | 0.104 | 50 |
| 5,6-DiHETrE | 337.20 > 145.00 | 52 | 16 | 10.06 | 0.250 | 0.083 | 100 |
| 13-HODE | 295.20 > 195.00 | 20 | 16 | 10.49 | 1.000 | 0.334 | 400 |
| 5-HEPE | 317.10 > 115.00 | 36 | 16 | 10.58 | 0.625 | 0.069 | 500 |
| 15,16 -EpODE | 293.10 > 235.00 | 34 | 16 | 10.91 | 0.500 | 0.083 | 200 |
| 15-HETE | 319.20 > 219.00 | 45 | 10 | 10.97 | 0.312 | 0.104 | 250 |
| 17,18-EpETE | 317.20 > 215.00 | 20 | 10 | 11.03 | 0.416 | 0.139 | 200 |
| 9,10-EpODE | 293.10 > 171.00 | 28 | 16 | 11.11 | 0.104 | 0.035 | 50 |
| 12,13-EpODE | 293.10 > 183.00 | 34 | 16 | 11.37 | 2.500 | 0.833 | 2000 |
| 11-HETE | 319.20 > 167.00 | 28 | 16 | 11.48 | 0.125 | 0.042 | 100 |
| 14,15-EpETE | 317.20 > 207.00 | 27 | 10 | 11.60 | 0.167 | 0.055 | 200 |
| 11,12-EpETE | 317.20 > 167.00 | 15 | 10 | 11.77 | 0.250 | 0.083 | 100 |
| 12-HETE | 319.10 > 179.00 | 44 | 16 | 11.82 | 0.500 | 0.167 | 400 |
| 8,9-EpETE | 317.20 > 127.00 | 20 | 16 | 11.99 | 1.250 | 0.416 | 200 |
| 8-HETE | 319.10 > 155.00 | 44 | 16 | 11.95 | 0.500 | 0.167 | 400 |
| 9-HETE | 319.00 > 151.00 | 28 | 16 | 12.26 | 1.250 | 0.139 | 500 |
| 5,6-EpETE | 317.20 > 189.00 | 20 | 10 | 12.35 | 0.500 | 0.167 | 400 |
| 5-HETE | 319.20 > 115.00 | 28 | 16 | 12.72 | 0.167 | 0.055 | 200 |
| 19,20-EpDPE | 343.20 > 299.00 | 27 | 10 | 12.80 | 0.625 | 0.208 | 250 |
| 12,13-EpOME | 295.20 > 195.00 | 20 | 16 | 13.02 | 0.312 | 0.104 | 250 |
| 14,15-EpETrE | 319.20 > 219.00 | 33 | 10 | 13.15 | 0.625 | 0.208 | 100 |
| 9,10-EpOME | 295.20 > 171.00 | 20 | 16 | 13.31 | 0.312 | 0.104 | 50 |
| 16,17 EpDPE | 343.20 > 274.00 | 20 | 10 | 13.40 | 1.250 | 0.416 | 500 |
| 13,14-EpDPE | 343.10 > 193.00 | 30 | 10 | 13.56 | 0.208 | 0.069 | 100 |
| 10,11-EpDPE | 343.10 > 153.00 | 36 | 10 | 13.73 | 0.250 | 0.028 | 40 |
| 7,8-EpDPE | 343.10 > 109.00 | 44 | 16 | 14.09 | 1.250 | 0.416 | 1000 |
| 8,9-EpETrE | 319.20 > 155.00 | 27 | 10 | 14.25 | 1.250 | 0.416 | 500 |
| 14,15 -EpEDE | 321.10 > 221.00 | 36 | 16 | 14.58 | 0.104 | 0.035 | 50 |
| 5,6-EpETrE | 319.20 > 191.00 | 20 | 10 | 14.61 | 2.500 | 0.832 | 1000 |
| 11,12-EpETrE | 319.20 > 167.00 | 27 | 10 | 14.93 | 0.208 | 0.069 | 250 |
| 8,9-EpEDE | 321.20 > 157.00 | 28 | 16 | 15.05 | 0.416 | 0.139 | 500 |
| 11,12-EpEDE | 321.20 > 181.00 | 36 | 16 | 15.14 | 0.416 | 0.139 | 500 |

### 2.2. Method validation

#### 2.2.1. LOQ and linearity

Limit of detection (LOD) and limit of quantification (LOQ) are essential parameters that indicate the lowest analyte concentrations that can be reliably measured by an analytical method. In this study, LOD is defined as the amount of a sample required to produce a signal-to-noise ratio (S/N) of 3 or greater, while the LOQ demands an S/N of 10 or higher to be acceptable. To determine these critical thresholds, four consecutive runs of calibration standards were analyzed within the same day. This approach allowed for establishment of LOQs ranging from 0.063 nM to 10 nM and LODs spanning from 0.021 nM to 5 nM, where the analytes were evaluated at a final volume of 100 μL ethanol (75% v/v with water containing a 10 nM concentration of CUDA which is the type II internal standard) (**Table 1**). The least favorable LOD and LOQ were observed for 12,13-DiHODE (0.416, 1.25 nM), 5,6-DiHETE (5, 10 nM), and 12,13-EpODE (0.833, 2.5 nM), whereas the best sensitivity or lowest LOD and LOQ belong to these three compounds 15,16-DiHODE, 14,15-DiHETE, and 9,10-DiHOME which both have LOD at 0.021 and LOQ at 0.063 nM) (**Table 1**). The linearity of the method was three orders of magnitude for most compounds when it is evaluated within the concentration range between the lower limit of quantification (LLOQ) and the upper limit of quantification (ULOQ), as determined by the calibration curves constructed for each analyte **(SI Figures 1, and 3)**.

#### 2.2.2. Assessing the accuracy and precision of the method

To evaluate the accuracy and precision of the method, we prepared five replicates of quality control samples including the lower limit of quantification (LLOQ), the lower quality control (LQC), middle-quality control (MQC), and higher quality control (HQC). For intra-day accuracy assessment, the percent difference between the mean concentration per analytical run and the expected concentration was determined. Five replicates were injected three times within the same day. This approach allowed for a robust evaluation of the method’s consistency and reliability during a single day of operation. Inter-day accuracy was also evaluated by injecting five replicates once daily for three consecutive days. This assessment provided insights into the method’s performance over multiple days, highlighting its stability and dependability across an extended period. Intra-day precision was evaluated as the relative standard deviation of three rounds of injection on the same day, while inter-day precision represented the relative standard deviation of the measurements (n=5, for each QC concentration) injected on three different days. Both precision values were calculated, and most of them were found to be within the acceptable criteria (≤ 25%) **(Table 2)**. Less than 20 compounds showed accuracy and precision out of the accepted range for at least one of the QCs, except for some of the LLOQ samples. The compounds that exhibited the least favorable accuracy and precision in the analysis were 5,6-DiHETE and 5,6-DiHETrE. This outcome was anticipated due to the compounds’ susceptibility to intramolecular lactonization **(SI Figure 4)**. ^30^ Furthermore, compounds such as 9-HODE, and 13-HODE displayed precision and accuracy values that fell outside the acceptable range. It is worth mentioning that the accuracy and precision of certain prostaglandins and lipoxins were also found to be lower than the acceptable criteria, which explains why these compounds were excluded from our analysis **(Table 2).** However, this observation does not raise concern, as these compounds are not naturally present in *C. elegans*. This is further supported by the fact that *C. elegans* lacks homologs for LOX and COX enzymes, which are responsible for the endogenous production of these compounds. ^31^ As a result, we expected that these compounds might not be produced in significant amounts. Furthermore, our oxylipin analysis with *C. elegans* also demonstrated that the metabolites from LOX and COX metabolic pathways were not detected in our method (**SI Table 6**). Therefore, the data related to these metabolites were excluded from this study; a list of these compounds is provided in supporting information (**SI Table 5**). The two figures of merit were acquired for all oxylipins listed in (**Table 2**), with the exception of metabolites from COX and LOX enzymes. Moving forward, we will primarily concentrate on CYP450 and EH enzyme metabolites. Despite these findings, in general, intra-day precision was found to be better than inter-day precision. Several factors could contribute to this observation, including instrument stability, reagent and sample stability, calibration curve stability, and instrument drift.

**Table 2.**
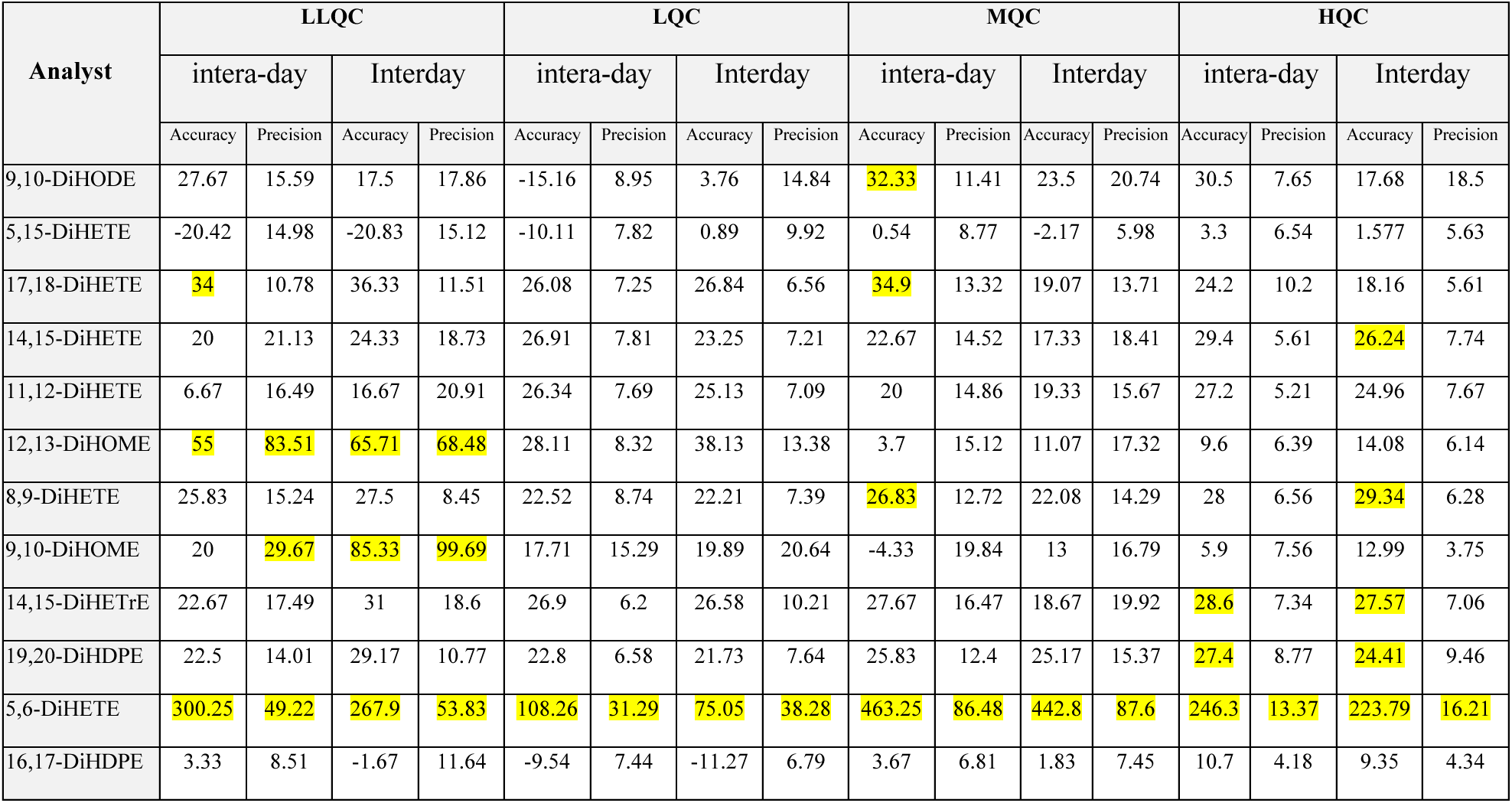
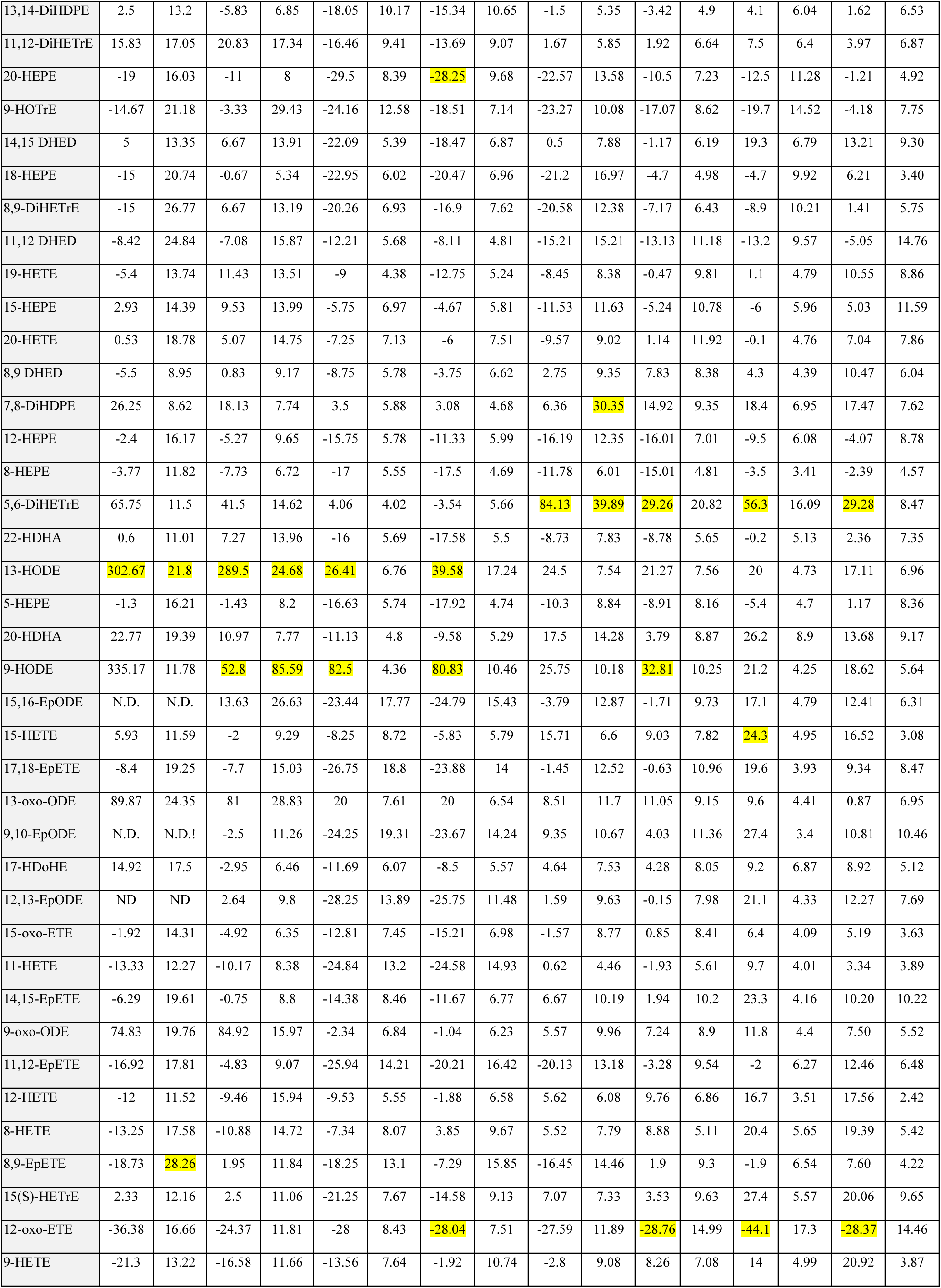
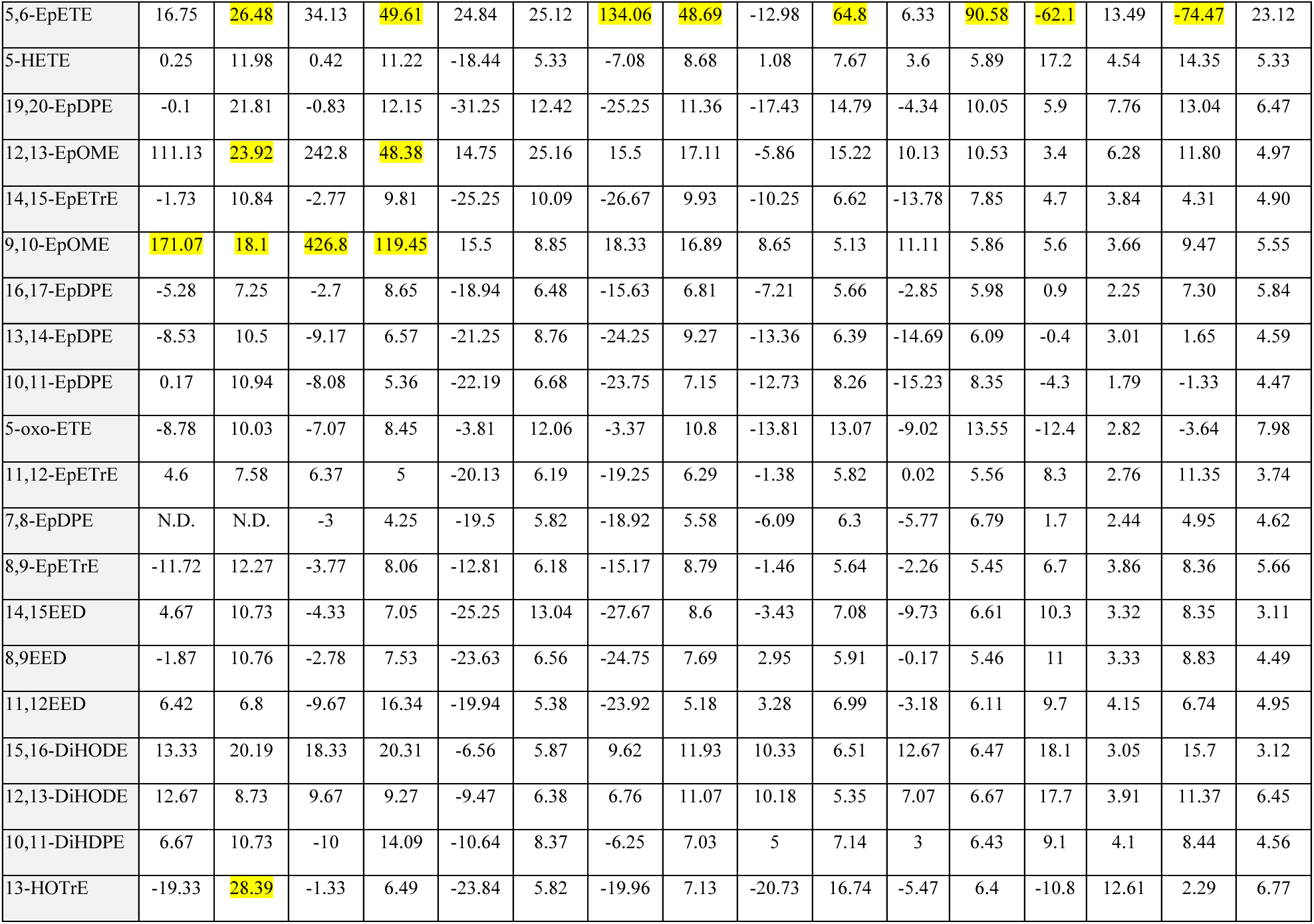
Accuracy and precision of oxylipins, Inter- and Intera-day precision represents the relative standard deviation (RSD) of the measurements (n=5). The intra-day accuracy was determined as the percent difference between the mean concentration per analytical run and the expected concentration. N.D. refers to the concentration below LOQ. Accuracy and precision of eicosanoids metabolites, Interday. precision represents the relative standard deviation of the measurements (n=5). The intra-day accuracy was determined as the percent difference between the mean concentration per analytical run and the expected concentration. The LLOQ, the LQC, MQC, and HQC concentration.

In this study, it is noteworthy to highlight that certain compound, including metabolites derived from DGLA, three distinct isomeric forms of EpEDE, and DHED, were successfully synthesized and comprehensively characterized for the very first time. As a result, the incorporation of these oxylipins in our method enables us to determine these important lipids in *C. elegans*, like DHED which induces neurodegeneration and ferroptosis, ^11^ thereby contributing valuable insights to the quantitative lipidomics and targeted metabolomics field.

#### 2.2.3 Evaluating recoveries and extraction efficiencies

Recoveries were calculated for QC samples, with results ranging from 72%-108% (n=5). These values indicate that most of the analytes were successfully extracted and recovered during the sample preparation process, suggesting a reliable and efficient methodology. The precision was observed to be satisfactory, with most RSDs falling below 12%. Our results demonstrate that the method provides consistent results across replicates, highlighting its suitability for accurate quantification of analytes. Moreover, the extraction efficiencies exhibited stability across different analytes, further supporting the method’s robustness and applicability to various compounds, the recovery result is comparable with other oxylipin studies in human plasma and serum (**SI Table 4).** ^32^

### 2.3. *C. elegance* sample analysis

To evaluate our developed method, we analyzed the oxylipins in *C. elegans* samples, where age-synchronized worms were grown on OP50, and collected on day 1 of adulthood (**SI Figure 5**).

#### 2.3.1. Optimizing the homogenization solvent

Tissue and animal samples, such as *C. elegans*, require homogenization prior to sample preparation to ensure efficient extraction of oxylipins. To achieve accurate quantification of PUFA metabolites in worm homogenates, it is essential to assess the solvent composition for optimal oxylipin recovery. ^33^ Considering this, two different solvents, phosphate-buffered saline (PBS) as a water-based solvent and isopropanol as an organic solvent, were tested in conjunction with bead-based homogenization for preparing *C. elegans* homogenates. A comparison of the two solvents revealed that a few metabolites demonstrated similar results for both PBS and isopropanol, with no statistically significant differences between them, such as 11,12 DiHETrE (**Figure 2A**). One potential reason for the lower efficiency of isopropanol as an organic solvent could be the formation of protein precipitates, which may interfere with the extraction process. Thus, the choice of homogenization solvent plays a critical role in the overall success of oxylipin extraction and quantification.

**Figure 2.**
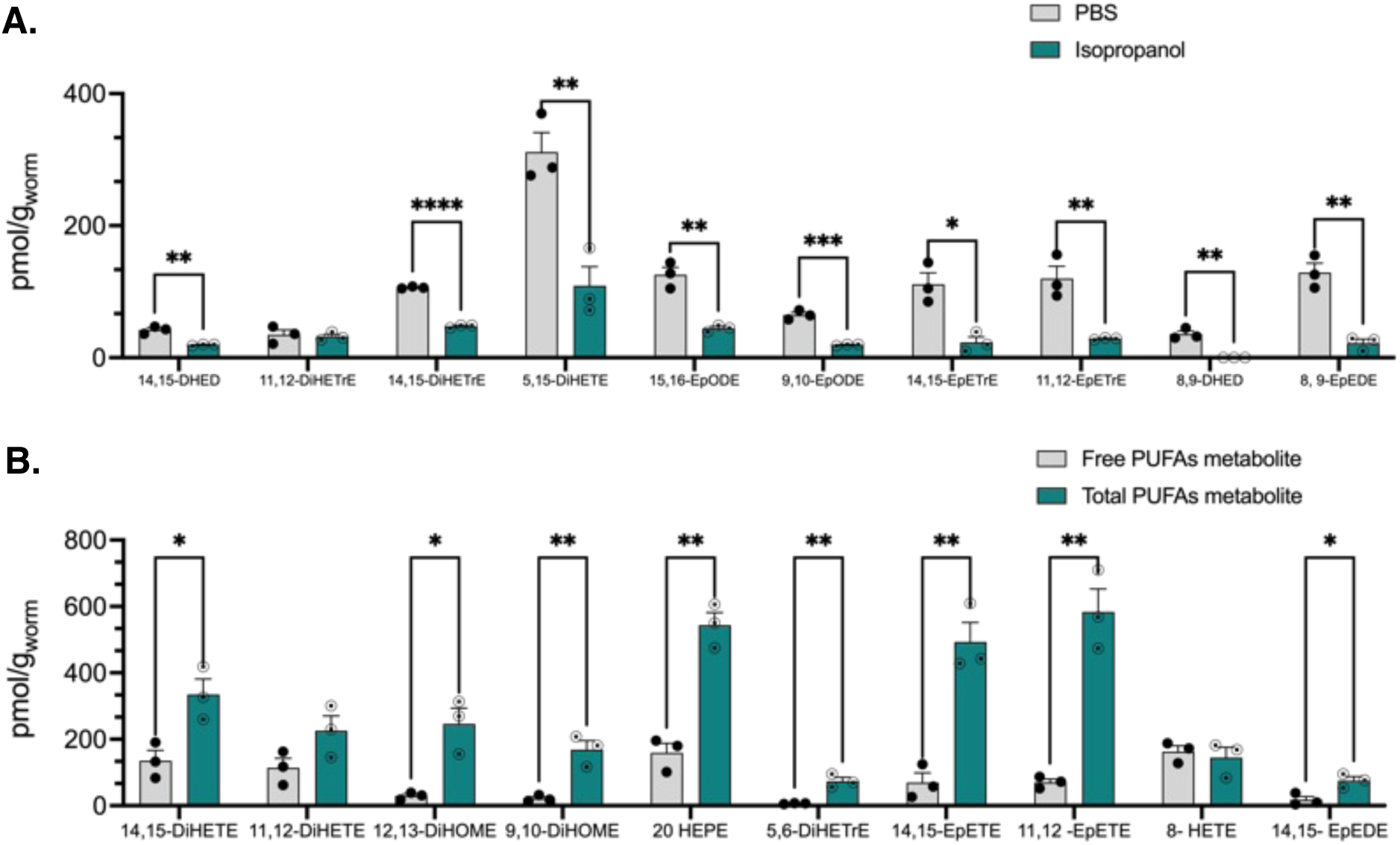
**A)** Optimization of solvent for C. elegans homogenization step, using two different solvents PBS and isopropanol. Oxylipin metabolites concentrations (pMol/g), mean ± SEM (n=3). Statistical differences between two different solvent groups were evaluated by multiple unpaired t-tests with *P ≤ 0.05, **P ≤ 0.01, ***P ≤ 0.001, ****P < 0.0001, non-significant is not shown). **B)** Oxylipin composition before and after hydrolysis in C. elegans samples. Control represents free fatty acids metabolites, whereas hydrolysis shows the total amount of oxylipin metabolites. Concentrations are mean ± SEM (n=3). Statistical differences between control and hydrolysis groups were evaluated by multiple unpaired t-tests with *P ≤ 0.05, **P ≤ 0.01, non-significant is not shown).

#### 2.3.2. Assessing oxylipin composition before and after hydrolysis in *C. elegans* sample

To gain a comprehensive understanding of the function of PUFA metabolites, ^34^ we needed to analyze both free and esterified oxylipins because studies have revealed that several oxylipins also initiate their effects through their phospholipid-esterified product. ^35^ In addition, recent studies revealed that the cell membrane can act as a reservoir for free epoxy-FAs, and an external stimulus can trigger the release of these esterified epoxy-FAs, triggering subsequent biological effects.^36^

To determine the total oxylipin levels, a well-established KOH/methanol method was used based on a published paper ^37^ (the detailed hydrolysis method is described in SI). After the hydrolysis process, certain oxylipins exhibited significantly different concentrations when compared to the nonhydrolyzed samples. These included both CYP450 metabolites, namely 11,12- EpETE, and 14,15- EpEDE, as well as epoxide hydrolase metabolites, specifically 9,10- DiHOME and 14,15- DiHETE.

However, for metabolites such as 8-HETE, and 11,12-DiHETE, the concentration difference between hydrolyzed samples and control samples was not statistically meaningful (**Figure 2B**). This observation underscores the importance of evaluating oxylipin composition both before and after hydrolysis in *C. elegans* samples. The variations in concentration might be attributed to the different susceptibilities of esterified oxylipins to hydrolysis or their varying affinities for enzymes involved in their metabolism. Further studies could provide insights into the specific factors contributing to these differences and improve our understanding of oxylipin metabolism in *C. elegans*.

To determine whether the alterations in oxylipin profile before and after hydrolysis were due to the breakdown of specific lipid metabolism or an alternation of the esterified oxylipin profiles within a membrane, a separate experiment was conducted by the identical hydrolysis using Type I IS only. During this experiment, we noticed that the relative abundance of all different internal standards under hydrolysis conditions is the same, which shows that the relative stability of all oxylipins under the hydrolysis condition is the same. Therefore, the significant increase in concentration of specific oxylipins after hydrolysis can be attributed to their relatively high level in the cell membrane as an esterified form.

#### 2.3.3. Matrix effect in *C. elegans* and ionization efficiency

Matrix effects (ME) pose significant challenges to quantitative LC-MS analysis by compromising the precision, sensitivity, and accuracy of the methodology. It is essential to thoroughly assess the presence of matrix effects during method development to ensure trustworthy analytical results ^38^. The matrix effect plays a significant role in oxylipin analysis in a variety of biological samples. Unlike other cases, it is not feasible to construct a calibration curve using the same matrix due to the endogenous oxylipins in most biological samples. Therefore, we were compelled to rely on 75% EtOH as the solvent to create our calibration curve. To quantitatively evaluate matrix effects in the post-extraction addition method, we will compare the concentration of the analyte in a standard solution with that of a post-extraction spiked with the analyte at an equal concentration. Our results indicated that compounds like PGB_2_-d_4_, 15(S)-HETE-d_8_, 9- HODE-d4, and 8,9-EET-d_11_ showed a significant matrix effect of more than 20%. This change indicates that the worm matrix decreases ionization efficiency in ESI, the calibration curve concentration, spiked concentration, and calculated matrix effect for each compound are shown in **Table 3**. An extracted ion chromatogram (XIC) of oxylipin in the calibration curve and day one worm samples is provided **(SI Figure 1).** Two common ways to tackle the ME are reducing the injection volume and diluting the sample before injection. None of these two strategies are applicable for oxylipin due to the extremely low concentration of these compounds in biological samples. We decided to normalize our result by the type I internal standard, this approach is the most appropriate technique available to decrease the ME.

**Table 3.** The matrix effect can be quantitatively evaluated by comparing the concentration of the analyte in standard solution (A) to that of a post-extract spiked with the analyte at the same concentration (B).

| Compound | Calibration curve (A) | Spiked (B) | Matrix effect % |
| --- | --- | --- | --- |
| PGB2-d4 | 15.63 | 19.73 | -26.25±5.56 |
| LTB4-d4 | 10.08 | 9.83 | 2.45± 8.28 |
| 8,9-DiHETrE-d11 | 15.70 | 18.83 | -19.96 ± 2.21 |
| 9-HODE-d4 | 16.70 | 20.63 | -23.55±9.22 |
| 15(S)-HETE-d8 | 20.50 | 23.30 | -13.66±5.22 |
| 5-HETE-d8 | 42.00 | 56.97 | -35.63± 7.49 |
| 8,9-EET-d11 | 42.10 | 56.50 | -34.19± 1.87 |

#### 2.3.4. Preparation of Age-synchronized worm

Traditional techniques for preventing progeny overgrowth during *C. elegans* lifespan and aging often employ 5-fluoro-2′-deoxyuridine (FUDR) for sterilization. While FUDR has been popularly employed to inhibit DNA and RNA synthesis, thereby eliminating progeny contamination, there are significant concerns tied to its use. ^39,40^ For example, the effects of FUDR have been primarily studied in wild-type (N2) worms. Notably, the interactions between FUDR and specific mutant strains have raised questions about its efficacy and reliability ^41,42^. Studies like those by Aitalhaj *et al.* and the *gas-1* strain underscore potential interferences, possibly skewing results and influencing longevity ^43,44^. Furthermore, the idea that the impact of FUDR sterilization is linear could be an oversimplification. External signals from the reproductive system have the potential to alter nematode lifespan ^45^, hinting at potential unexplored dimensions of FUDR effects. Additionally, our previous study raised concerns over the possible interference of FUDR with CYP-EH metabolites

Amid the aforementioned challenges with FUDR, our approach transitioned to a filtration method to achieve age synchronization. ^46^ This method was meticulously designed to draw a clear boundary between adult worms and their progeny, resulting in a striking separation efficiency of over 99% **(SI Figure 5).** It is noteworthy to mention that while there was a minor loss of adults during filtration, primarily because live animals occasionally adhered to the filter pores or passed through during the washing process, this loss was quantified to be less than 15% **(SI Figure 5).** The high efficacy of this method became even more pronounced as the worms aged. Initial stages required daily filtrations, but as the adult worms advanced in their life cycle and stopped producing progeny, the filtration frequency diminished. After the initial week, the predominant aim shifted from rigorous progeny segregation to primarily transferring worms to plates with fresh food.

#### 2.3.5. Analysis of *C. elegans* with and without EHI

To determine whether our method can quantify the oxylipin changes in *C. elegans*, we quantified the oxylipin profiles in *C. elegans* with and without the treatment with an epoxide hydrolase inhibitor (EHI). Harris *et al.* showed that 12-(3-((3s,5s,7s)-adamantan-1-yl) ureido) dodecanoic acid (AUDA), blocks the conversion of epoxyoctadecenoic acids to dihydroxyoctadecenoic acid by inhibiting *C. elegans* epoxide hydrolases. ^47^

The oxylipin profile of AUDA-treated worm as compared with vehicle control revealed an overall increase in the levels of epoxy-FA metabolites, and a decrease in dihydroxy-FA metabolites, with the most pronounced change observed in (i) 12,13-DiHOME among LA metabolites, (ii) 8,9-DHED and 14,15-DHED for DGLA metabolites, (iii)11,12-EpETrE, and 14, 15-DiHETrE for AA metabolites, and (iv) 17,18-EpETE, 11,12-DiHETE, and 14,15-DiHETE for EPA metabolites (**Figure 3**). This observation is intriguing, as it implies that the connection between epoxide hydrolase activity and overall oxylipin levels could be more complex than initially thought. In particular, the presumption is that inhibiting epoxide hydrolase activity would stabilize or increase the endogenous level of epoxy fatty acids and decrease the corresponding epoxide hydrolase products (dihydroxy fatty acids). This observation could be attributed to the presence of alternative pathways involved in the metabolism of epoxy- and dihydroxy-FA metabolites, as well as potential feedback regulation of other enzymes that play a part in this process. In addition, the diverse effects caused by the supplementation of AUDA on a variety of epoxy-FA-to-dihydroxyPUFA ratios could also be attributed to the selectivity of the *C. elegans* EHs. Moreover, we found a higher concentration of different EPA metabolites, and the lower concentration of the ALA metabolites compared to the other PUFA metabolites. This pattern aligns with the levels of their respective parent PUFAs found in the worm. ^48^

**Figure 3.**
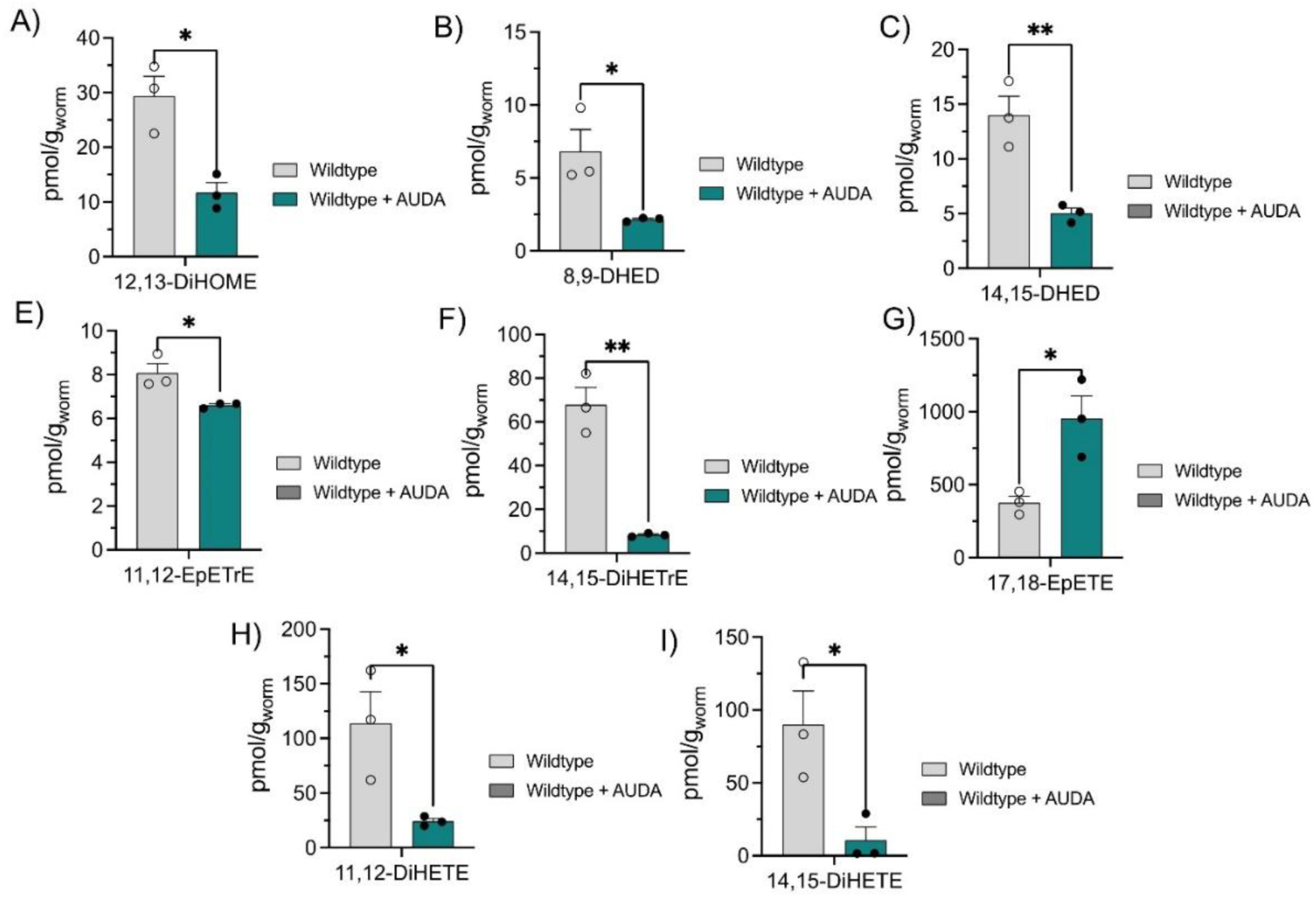
PUFA metabolites of C. *elegans* in the first day of adulthood, with and without epoxide hydrolase inhibitor treatment. The concentration of each compound (pmol/g) represents a replicate of independent populations of whole worm lysate that were 10 mg or greater, mean ± SEM (n=3). Statistical differences between two different groups were evaluated by multiple unpaired t-tests with *P ≤ 0.05, **P ≤ 0.01, and non-significant is not shown).

It is worth mentioning that interpreting the effect of EH inhibitors could not be done correctly by solely tracking the Ep-PUFAs or dihydroxy-FA levels independently, as there is a physiological balance between these different oxylipins (**Figure 1**). Therefore, to further explore the main effect of EH inhibition, the epoxy-to-dihydroxy ratio of different PUFA oxylipin metabolites was studied, which is generally considered a marker of EH inhibition *in vivo. ^49,50^*Our results show that wildtype worms treated with AUDA exhibit an overall increase in the epoxy-to-dihydroxy fatty acid ratio, with the most significant increase related to the 11,12- (EED/DHED), 14,15-(EED/DHED), 11,12-(EEQ/DiHETE), and 17,18-(EEQ/DiHETE). This corroborates with previous studies that reported an increase in the epoxy-to-dihydroxy fatty acid ratio upon administration of EHI (**Figure 4A-D**).

**Figure 4.**
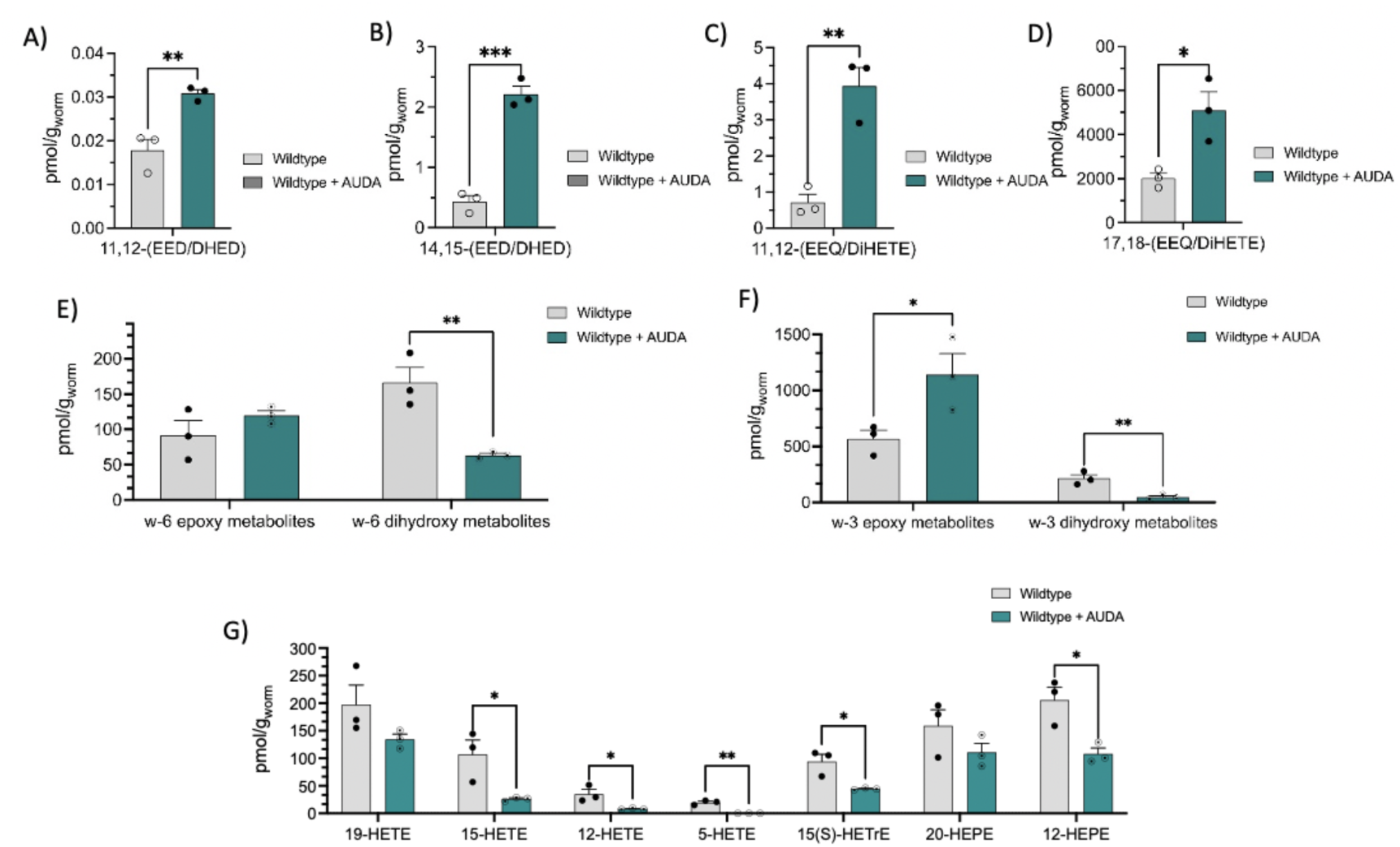
**(A-D)** The ratio of epoxy to dihydroxy PUFAs metabolite of C. elegans in the first day of adulthood, with and without epoxide hydrolase inhibitor treatment. The concentration of each compound (pMol/g) represents a replicate of independent populations of whole worm lysate that were 10 mg or greater, mean ± SEM (n=3). Statistical differences between two different groups were evaluated by multiple unpaired t-tests with *P ≤ 0.05, **P ≤ 0.01, ***P ≤ 0.001, and non-significant is not shown). **(E -G).** Represent the comparison of the total, ω-6, and ω-3 PUFA epoxy, dihydroxy metabolites and hydroxy metabolites in C. elegans on the first day of adulthood, with and without epoxide hydrolase inhibitor treatment. The concentration of each compound (pmol/g) represents a triplicate of independent populations of whole worm lysate that were 10 mg or greater, mean ± SEM (n=3). Statistical differences between two different groups were evaluated by multiple unpaired t-tests with *P ≤ 0.05, **P ≤ 0.01, and non-significant is not shown).

We also compared the overall level of CYP-EH metabolites related to ω-3 and ω-6 PUFAs (**Figure 4 E, and** **F**). The wildtype worms show a higher level of ω-3 Ep-PUFAs (600-800 pmol/g) compared to ω-6 Ep-PUFAs (90-120 pmol/g), while the dihydroxy-FA level was almost similar for both ω-3 and ω-6-PUFAs (150-200 pmol/g). Intriguingly, administration of AUDA resulted in a significant increase in ω-3 Ep-PUFAs and a significant decrease in ω-3 dihydroxy- FAa. On the other hand, we only observed a decrease in ω-6 Dihydroxy-FAs, while the ω-6 Ep-PUFAs remained unchanged.

Besides, we could measure several hydroxy-PUFA in wild-type worms (**Figure 4G**). It should be noted that the hydroxy-PUFAs detected in this research are predominantly generated via three main pathways in mammals: (i) LOX enzymes, (ii) non-enzymatic oxidation, and (iii) cytochrome P450 (CYP450) enzymes. Given the absence of LOX homologs in *C. elegans*, the production of hydroxy-PUFAs in this organism is likely primarily facilitated by non-enzymatic oxidation, CYP450 pathways, or an undiscovered enzyme with monohydroxylation activity.^51-53^ This observation highlights the unique metabolic processes in *C. elegans* and the potential for further study of these mechanisms. We also found that AUDA treatment could change the hydroxy-PUFA levels in worms, with a statistically significant decrease in 5-HETE, 12-HETE, and 15-HETE. Whereas for the hydroxy metabolites like 19-HETE and 20-HEPE, the difference caused by AUDA treatment was not statistically significant. These findings emphasize the complex effects of AUDA treatment on hydroxy-PUFA levels and point to possible indirect influences on other enzymatic pathways or feedback regulation mechanisms. Further research is necessary to determine the potential roles of these changes and to elucidate the precise mechanisms underlying the observed alterations in hydroxy-PUFA levels.

## 3. Conclusion

Overall, these findings highlight the efficacy of our method for quantifying the oxylipin profile in *C. elegans* and offer valuable insights into the effects of epoxide hydrolase inhibition on PUFA metabolites. The method could be used to further reveal intricate relationships between enzymatic activity, oxylipin levels, and the roles of different metabolic pathways in *C. elegans*, pointing to the possibility of complex regulatory mechanisms at play. By providing a comprehensive picture of alterations in oxylipin profiles after AUDA treatment, our work lays the foundation for future investigations into the underlying mechanisms and potential therapeutic applications of epoxide hydrolase inhibitors.

This study aimed to develop a reliable and accurate LC-MS/MS method for quantifying the oxylipin profile in the *C. elegans* animal model. The developed method can be employed in future studies to investigate the effects of various diseases, treatments, or genetic manipulations on the oxylipin profile and to assess potential therapeutic interventions using *C. elegans* as an animal model. Ultimately, the knowledge gained from these studies may contribute to a better understanding of the role of oxylipins in health and disease and may potentially lead to the development of novel therapeutic strategies. By further refining the method and expanding the range of oxylipins analyzed, researchers can continue to build upon the foundation established in this study and enhance our understanding of the complex interplay between oxylipin metabolism and various biological processes.

## 4. Acknowledgment

This study was supported, by the National Institute of Aging R03 AG075465, the Pearl Aldrich Endowment for Aging Research, and startup funding from Michigan State University. The Peal Aldrich Endowment for Aging Research Student award partially supported. We acknowledge the MSU RTSF Mass Spectrometry and Metabolomics Core Facilities for support with oxylipin and lipidomic analysis. We would like to thank Prof. Daniel Jones for his advice help and assistance. Prism and Biorender partly generate figures.

## Notes

### Competing Interest Statement

The authors have declared no competing interest.

