## supplemental Information for "Quantitative Profiling Method for Oxylipins in Neurodegenerative Diseases by Liquid Chromatography Coupled with Tandem Mass Spectrometry"

### Supporting Information

#### Experimental and methods

##### Chemicals

1) The following compounds were all purchased from Cayman Chemical Company (Ann Arbor, MI):

8,9-Dihydroxyeicosatetraenoic acid (8,9-DiHETE), (±)7,8-dihydroxydocosa-pentaenoic acid (7,8 DiHDPE); 19,20 DiHDPE, 16,17 DiHDPE, 13,14 DiHDPE, 10,11 DiHDPE, (±)7,8-epoxy docosapentaenoic acid (7,8 EpDPE), 13,14 EpDPE, 10,11 EpDPE, (±)5,6-dihydroxy-eicosatrienoic acid (5,6 DiHETrE), (±)5,6-epoxyeicosatrienoic acid (5,6 EpETrE) , 11,12-dihydroxyeicosatetraenoic acid (11,12 DiHETE), 5-oxo-eicosatetraenoic acid(5 Oxo ETE), (±)9-hydroxy-10E,12Z-octadecadienoic acid (9 HODE), 13-keto-9Z,11E- octadecadienoic acid (13 oxo ODE), 9-oxo-octadecadienoic acid (9 oxo ODE), (±)5-hydroxyeicosatetraenoic acid (5HETE); 20 HETE, 11 HETE , 9 HETE, 12 HETE; (±)8 HETE, (±)17-hydroxydocosahexaenoic acid (17 HDoHE), 5S,15S-dihydroxyeicosatetraenoic acid (5,15 DiHETE), 8S,15S-dihydroxyeicosatetraenoic acid (8,15 DiHETE), (±)-12-hydroxyeicosapentaenoic acid (12 HEPE); 15 HEPE, 5 HEPE , 8 HEPE,12-oxo-eicosatetraenoic acid (12-oxo-ETE), 13S-hydroxy-octadecatrienoic acid (13-HOTrE); 15(S)HETrE, 5S,12R-dihydroxyeicosatetraene-1,20-dioic acid (20-COOH-LTB4),5S,12R,20-trihydroxy-6Z,8E,10E,14Z-eicosatetraenoic acid. (20-OH-LTB4) 9S-hydroxyoctadecatrienoic acid (9(s)HOTrE); 12,13-epoxy-9-keto-10(trans)-Octadecenoic Acid (trans-EKODE-(E)-lb),Leukotriene-B3 (LTB3); LTB5, Leukotriene-E4 (LTE4), Prostaglandin-B2 (PGB2), Prostaglandin-D1 (PGD1),PGD3, Prostaglandin-E1 (PGE1), PGE3, Prostaglandin-J2(PGJ2), Resolvin-E1 Prostaglandin-F2α (PGF2a), Thromboxane-B2-d4 (TXB2-d4)Leukotriene-B4-d4 (LTB4-d4), 9S-hydroxy-10E,12Z-octadecadienoic-9,10,12,13-d4 acid- d4 (9 HODE-d4), 6- oxo-9S,11R,15S-trihydroxy-13E-prostenoic acid (6 keto PGF1α) .

2) The following compounds were received as a kind gift from the MSU Metabolomic facility :Thromboxane B2 (TXB2), lipoxin A4 (LXA4), 17,18DiHETE, 8,9 EpETrE, 17,18 EpETE, 14,15 DiHETrE , LTB4, 11,12 EpETE, 8-iso-PGF2a, 15 oxo ETE, 10,11 EpDPE, 19,20 EpDPE, 11,12 EpETrE, PGD2,docosahexaenoic acid (DHA), eicosapentaenoic acid (EPA), arachidonic acid (AA), 12,13-dihydroxyoctadeca-9,15-dienoic acid (12,13DiHODE); 9,10 DiHODE, 15,16 DiHODE, acid, 1-cyclohexyl-dodecanoic acid urea (CUDA), 12,13-dihydroxyoctadecenoic acid (12,13 DiHOME); 9,10 DiHOME, 9,10 EpODE, 15,16 EpODE, 14, 15EpETrE, standards, 6-keto-PGF1a-d4, 5-HETE-d8, 8,9-EET-d11, AA-d8, 15(S)-HETE-d8, PGB2-d4, 8,9-DiHETrE-d11, 9-HODE-d4, LTB4-d4, PGE2-d9.

3) These compounds were synthesized in-house, and the synthetic procedures are described in For characterization details, please see our previous publication[1]:

14,15-Epoxyeicosadienoic acid (14,15 EED); 8,9 EED, 11,12-EED, 14,15-di-hydroxyeicosadienoic acids (14,15 DHED);8,9 DHED, 11,12 DHED.

Phosphate buffered saline (PBS) was purchased from Gibco, (Grand Island, NY) ethanol, isopropanol, methanol, ethyl acetate, acetonitrile, and glacial acetic acid (HPLC Grade) were purchased from Fisher Scientific (Pittsburgh, PA, USA). Oasis HLB 60 mg SPE cartridges were purchased from Waters Co. (Milford, MA)

**Internal standards**

To determine the recovery of extraction and instrumental variation, two different types of compounds were used as internal standards. The following deuterated compounds were used as Type I internal standards: 6-keto-PGF1a-d4, 5-HETE-d8, 8,9-EET-d11, AA-d8, 15(S)-HETE-d8,

PGB2-d4, 8,9-DiHETrE-d11, 9-HODE-d4, LTB4-d4, PGE2-d9. The type I internal standards were added to the samples before the SPE. The physical and chemical properties similarity between Type I internal standard and targeted PUFA metabolites helps to calculate the extraction recovery of prostaglandins, diols, epoxides, and other oxylipins. In this regard, the analytes were linked to their corresponding Type I internal standards, based on their retention time, for the purpose of quantification.

A type II internal standard was added at the last step before injection to LC-MS/MS, to normalize changes in volume and other instrumental variations. A synthetic compound, CUDA which is a fatty acid metabolite mimic, was used as the type II internal standard (10 nM in 75% EtOH) **(SI- Table 1).**

**Standard curve preparation**

A 12-step dilution from the original stock of a mixture of PUFA metabolites with known concentrations was used to generate 10 concentration calibration standards in EtOH with a constant concentration of type I **(SI Table 2),** and type II CUDA (10 nM). Another calibration curve was made for the calculation of the extraction recovery purposes with different concentrations of type I and constant concentrations of type II (10nM). These calibration standards were kept in an amber vial (Fisher Scientific), sealed under argon gas, and stored at −80°C (figure 3 A). Curves were plotted using the response of the analyte, which is the ratio of the analyte area to its internal standard area, against the concentration with 1/x weighting factors in the regression. Regression analysis yielded $R^2$ values of 0.998 or greater for each analyte.

**SI Table 1.** Type I IS the name and concentration in the stock solution and in the calibration curve.

| Compound | Function | IS concentration stock (nM) | Concentration in calibration curve (nM) |
|---|---|---|---|
| 6-keto-PGF1a-d4 | 10 | 1000 | 40 |
| 5-HETE-d8 | 6 | 1000 | 40 |
| 8,9-EpETrE-d11 | 5 | 1000 | 40 |
| AA-d8 | 1 | 1000 | 40 |
| 15(S)-HETE-d8 | 4 | 500 | 20 |
| PGB2-d4 | 7 | 400 | 16 |
| 8,9-DiHETrE-d11 | 2 | 400 | 16 |
| 9-HODE-d4 | 3 | 400 | 16 |
| LTB4-d4 | 8 | 250 | 10 |
| PGE2-d9 | 9 | 100 | 4 |

**SI Table 2.** Different concentration categories of PUFAs metabolite (A-I) based on the sensitivity of the instrument and biological level of oxylipin. (12 step-dilution in the calibration curve)

| Conc. A (nM) | Conc. B (nM) | Conc. C (nM) | Conc. D (nM) | Conc. E (nM) | Conc. F (nM) | Conc. G (nM) | Conc. H (nM) | Conc. I (nM) |
|---|---|---|---|---|---|---|---|---|
| 0.09 | 0.04 | 0.02 | 0.018 | 0.011 | 0.008 | 0.005 | 0.002 | 0.0005 |
| 0.27 | 0.14 | 0.07 | 0.05 | 0.034 | 0.026 | 0.0138 | 0.007 | 0.0014 |
| 0.83 | 0.41 | 0.21 | 0.17 | 0.104 | 0.08 | 0.041 | 0.021 | 0.0041 |
| 2.5 | 1.25 | 0.625 | 0.5 | 0.3125 | 0.25 | 0.125 | 0.0625 | 0.0125 |
| 5 | 2.5 | 1.25 | 1 | 0.625 | 0.5 | 0.25 | 0.125 | 0.025 |
| 10 | 5 | 2.5 | 2 | 1.25 | 1 | 0.5 | 0.25 | 0.05 |
| 20 | 10 | 5 | 4 | 2.5 | 2 | 1 | 0.5 | 0.1 |
| 40 | 20 | 10 | 8 | 5 | 4 | 2 | 1 | 0.2 |
| 80 | 40 | 20 | 16 | 10 | 8 | 4 | 2 | 0.4 |
| 200 | 100 | 50 | 40 | 25 | 20 | 10 | 5 | 1 |
| 400 | 200 | 100 | 80 | 50 | 40 | 20 | 10 | 2 |
| 1000 | 500 | 250 | 200 | 125 | 100 | 50 | 25 | 5 |
| 2000 | 1000 | 500 | 400 | 250 | 200 | 100 | 50 | 10 |

**SI Table 3.** Linear gradient chromatographic method. The mobile phase comprised 0.1% acetic acid in water (mobile phase A) and ACN (84%) and MeOH(16%)(mobile phase B), in a total run time of 21 min

| Time | %Aqueous phase (A) | %Organic phase (B) | Flow rate (ml/min) |
|---|---|---|---|
| initial | 65 | 35 | 0.25 |
| 1 | 60 | 40 | 0.25 |
| 3 | 45 | 55 | 0.25 |
| 8.5 | 35 | 65 | 0.25 |
| 12.5 | 28 | 72 | 0.25 |
| 15 | 18 | 82 | 0.25 |
| 16 | 0 | 100 | 0.25 |
| 18.1 | 65 | 35 | 0.25 |

**SI Table 4.** Calculated recovery percentage based on type I standard to study the efficiency of solid phase extraction, n=3

| Internal standard | Spiked conc. (nM) | RC % |
|---|---|---|
| **6 keto PGF1 a-d4** | 40 | 72.66± 7.3 |
| **PGE2-d9** | 4 | 95.35±7.8 |
| **PGB2-d4** | 16 | 101.38±6.2 |
| **LTB4-d4** | 10 | 81.84±4.8 |
| **8,9-DiHETrE-d11** | 16 | 108.71±3.9 |
| **9-HODE-d4** | 16 | 86.47±2.5 |
| **15-HETE-d8** | 20 | 96.23±4.4 |
| **5-HETE-d8** | 40 | 95.3±3.9 |
| **8,9-EET-d11** | 40 | 76.48±11.5 |

**SI Table 5.** The list of compounds that were excluded from the paper because they were COX or LOX enzymes' metabolites. Some compounds were excluded because the accuracy and precision were not in an acceptable range.

| Analytes | MRM | Cone Volt. | Collision Volt. | RT (min) | LLOQ (nM) | LOD (nM) | ULOQ (nM) |
|---|---|---|---|---|---|---|---|
| 20-COOH-LTB4 | 365.10 > 195.00 | 52 | 22 | 3.08 | 1.25 | 0.416 | 1000 |
| 6-Keto-PGF1a | 369.30 > 163.00 | 21 | 22 | 3.22 | 0.625 | 0.208 | 500 |
| 20-OH-LTB4 | 351.10> 195.00 | 20 | 16 | 3.25 | 0.250 | 0.125 | 100 |
| 8-iso-PGF2a | 353.20 > 193.00 | 44 | 28 | 3.79 | 0.312 | 0.104 | 250 |
| PGE3 | 349.10 > 269.00 | 20 | 16 | 3.82 | 0.208 | 0.069 | 500 |
| TXB2 | 369.20 > 169.00 | 51 | 16 | 3.85 | 0.625 | 0.208 | 500 |
| PGD3 | 349.20 > 269.00 | 28 | 16 | 4.00 | 0.208 | 0.069 | 500 |
| PGF2a | 353.10 > 193.00 | 52 | 22 | 4.23 | 0.250 | 0.083 | 200 |
| PGE2 | 351.10 > 271.00 | 44 | 16 | 4.32 | 0.125 | 0.042 | 100 |
| PGE1 | 353.20 > 223.00 | 28 | 28 | 4.43 | 1.250 | 0.416 | 500 |
| RV D2 | 375.20 > 175.00 | 28 | 22 | 4.44 | 0.625 | 0.312 | 250 |
| PGD1 | 353.10 > 205.00 | 28 | 28 | 4.5 | 1.250 | 0.416 | 200 |
| PGD2 | 351.20 > 271.00 | 20 | 16 | 4.53 | 0.250 | 0.083 | 100 |
| LTD4 | 495.30 > 177.00 | 20 | 16 | 4.86 | 0.625 | 0.208 | 500 |
| LXA4 | 351.20 > 115.00 | 55 | 16 | 4.90 | 0.312 | 0.104 | 250 |
| PGB2 | 333.10 > 175.00 | 44 | 22 | 5.74 | 0.312 | 0.104 | 250 |
| LTB5 | 333.10 > 195.00 | 44 | 16 | 5.78 | 0.250 | 0.083 | 200 |
| LTE4 | 438.10 > 333.00 | 36 | 16 | 5.87 | 1.250 | 0.416 | 500 |
| 6-trans-LTB4 | 335.10 > 195.00 | 28 | 16 | 6.66 | 0.250 | 0.083 | 40 |
| LTB4 | 335.20 > 195.00 | 21 | 16 | 6.97 | 0.250 | 0.083 | 200 |
| LTB3 | 337.20 > 195.00 | 60 | 16 | 8.44 | 0.250 | 0.083 | 200 |
| EKODE | 309.10 > 209.00 | 20 | 10 | 8.90 | 0.500 | 0.167 | 200 |
| 15-deoxy-PGJ2 | 315.20 > 271.00 | 39 | 10 | 9.35 | 0.083 | 0.028 | 100 |
| 13-oxo-ODE | 293.20 > 113.00 | 15 | 22 | 11.08 | 0.312 | 0.104 | 250 |
| 17-HDoHE | 343.20 > 281.00 | 28 | 10 | 11.13 | 1.250 | 0.416 | 1000 |
| 15-oxo-ETE | 317.20 > 113.00 | 45 | 16 | 11.45 | 0.083 | 0.028 | 200 |
| 9-oxo-ODE | 293.20 > 185.00 | 27 | 16 | 11.63 | 0.500 | 0.167 | 200 |
| 12-oxo-ETE | 317.10 > 273.00 | 28 | 16 | 12.20 | 0.416 | 0.139 | 200 |
| 5-oxo-ETE | 317.20 > 203.00 | 20 | 16 | 13.80 | 1.250 | 0.139 | 500 |

**SI Table 6.** The oxylipin profiles of *C. elegans* at day 1 adult that is either treated with AUDA or vehicle.

| Compound | Control 1 (pmol/gworm) | Control 2 (pmol/gworm) | Control 3 (pmol/gworm) | AUDA 1 (pmol/gworm) | AUDA 2 (pmol/gworm) | AUDA 3 (pmol/gworm) |
|---|---|---|---|---|---|---|
| 20-COOH-LTB4 | LOD> | LOD> | LOD> | LOD> | LOD> | LOD> |
| 6-Keto-PGF1a | LOD> | LOD> | LOD> | LOD> | LOD> | LOD> |
| 20-OH-LTB4 | LOD> | LOD> | LOD> | LOD> | LOD> | LOD> |
| 8-iso-PGF2a | LOD> | LOD> | LOD> | LOD> | LOD> | LOD> |
| PGE3 | LOD> | LOD> | LOD> | LOD> | LOD> | LOD> |
| TXB2 | LOD> | LOD> | LOD> | LOD> | LOD> | LOD> |
| PGD3 | LOD> | LOD> | LOD> | LOD> | LOD> | LOD> |
| PGF2a | LOD> | LOD> | LOD> | LOD> | LOD> | LOD> |
| PGE2 | LOD> | LOD> | LOD> | LOD> | LOD> | LOD> |
| PGE1 | LOD> | LOD> | LOD> | LOD> | LOD> | LOD> |
| RV D2 | LOD> | LOD> | LOD> | LOD> | LOD> | LOD> |
| PGD1 | LOD> | LOD> | LOD> | LOD> | LOD> | LOD> |
| PGD2 | LOD> | LOD> | LOD> | LOD> | LOD> | LOD> |
| LTD4 | LOD> | LOD> | LOD> | LOD> | LOD> | LOD> |
| LXA4 | LOD> | LOD> | LOD> | LOD> | LOD> | LOD> |
| PGB2 | LOD> | LOD> | LOD> | LOD> | LOD> | LOD> |
| LTB5 | LOD> | LOD> | LOD> | LOD> | LOD> | LOD> |
| LTE4 | LOD> | LOD> | LOD> | LOD> | LOD> | LOD> |
| 6-trans-LTB4 | LOD> | LOD> | LOD> | LOD> | LOD> | LOD> |
| LTB4 | LOD> | LOD> | LOD> | LOD> | LOD> | LOD> |
| LTB3 | LOD> | LOD> | LOD> | LOD> | LOD> | LOD> |
| EKODE | LOD> | LOD> | LOD> | LOD> | LOD> | LOD> |
| 15-deoxy-PGJ2 | LOD> | LOD> | LOD> | LOD> | LOD> | LOD> |
| 9,10-EpOME | 13.96 | 42.42 | 25.07 | 54.22 | 47.50 | 47.26 |
| 9,10-DiHOME | 32.50 | 14.55 | 18.06 | 23.33 | 26.33 | 23.71 |
| 8,9- EEDs | 0.19 | 0.19 | 0.19 | 20.89 | 20.50 | 18.55 |
| 8,9-DHED | 9.81 | 5.45 | 5.22 | 2.22 | 2.00 | 2.26 |
| 5,6 -EpETrE | 173.08 | 109.09 | 165.67 | 0.38 | 0.38 | 0.38 |
| 5,6-DiHETrE | 7.88 | 7.27 | 5.22 | 7.78 | 7.00 | 0.38 |
| 15(16)-EpODE | 0.04 | 0.04 | 0.04 | 0.04 | 0.04 | 0.04 |
| 15,16-DiHODE | 4.62 | 5.15 | 2.99 | 3.78 | 2.33 | 2.74 |
| 12, 13- EpOME | 44.42 | 67.88 | 10.60 | 30.00 | 28.67 | 22.42 |
| 12,13-DiHOME | 30.77 | 34.79 | 22.51 | 15.11 | 11.17 | 8.87 |
| 14,15-EEDs | 9.62 | 3.80 | 4.48 | 11.78 | 10.33 | 10.97 |
| 14,15-DHED | 17.12 | 13.76 | 11.10 | 5.78 | 4.17 | 5.16 |
| 11,12-EED | 0.19 | 0.19 | 0.19 | 0.19 | 0.19 | 0.19 |

| | | | | | | |
|---|---|---|---|---|---|---|
| **11,12-DHED** | 14.81 | 9.09 | 9.25 | 6.44 | 5.83 | 5.97 |
| **14,15 -EpETrE** | 14.42 | 6.36 | 7.76 | 8.67 | 4.83 | 2.42 |
| **14,15-DiHETrE** | 82.12 | 66.52 | 55.07 | 9.11 | 8.33 | 7.58 |
| **11,12-EpETrE** | 7.69 | 7.58 | 8.96 | 6.67 | 6.67 | 6.45 |
| **11,12-DiHETrE** | 16.54 | 5.76 | 11.34 | 2.44 | 2.50 | 2.58 |
| ***17(18)-EpETE*** | 453.46 | 381.01 | 297.31 | 1221.11 | 690.50 | 951.61 |
| ***17,18-DiHETE*** | 0.19 | 0.19 | 0.19 | 0.19 | 0.19 | 0.19 |
| ***14,15 -EpETE*** | 57.50 | 124.55 | 26.57 | 81.78 | 45.67 | 49.03 |
| ***14,15-DiHETE*** | 53.38 | 83.33 | 132.84 | 28.89 | 1.67 | 1.61 |
| ***11,12 -EpETE*** | 86.54 | 72.27 | 52.09 | 128.00 | 69.33 | 89.35 |
| ***11,12-DiHETE*** | 162.31 | 62.12 | 117.16 | 28.67 | 23.83 | 20.16 |
| ***8,9 -EpETE*** | 74.42 | 34.24 | 41.49 | 44.67 | 20.00 | 30.32 |
| ***8,9-DiHETE*** | 38.08 | 15.76 | 27.91 | 11.33 | 12.33 | 9.03 |
| ***5,6-EpETE*** | 1116.73 | 835.30 | 806.12 | 1448.67 | 643.17 | 1071.77 |
| ***5,6-DiHETE*** | 517.12 | 319.09 | 538.81 | 404.00 | 433.17 | 305.65 |
| **19(20)-EpDPE** | LOD> | LOD> | LOD> | LOD> | LOD> | LOD> |
| **16(17)-EpDPE** | LOD> | LOD> | LOD> | LOD> | LOD> | LOD> |
| **13(14)-EpDPE** | LOD> | LOD> | LOD> | LOD> | LOD> | LOD> |
| **10(11)-EpDPE** | LOD> | LOD> | LOD> | LOD> | LOD> | LOD> |
| **7(8)-EpDPE** | LOD> | LOD> | LOD> | LOD> | LOD> | LOD> |
| **19,20-DiHDPE** | LOD> | LOD> | LOD> | LOD> | LOD> | LOD> |
| **16,17-DiHDPE** | LOD> | LOD> | LOD> | LOD> | LOD> | LOD> |
| **13,14-DiHDPE** | LOD> | LOD> | LOD> | LOD> | LOD> | LOD> |
| **10,11-DiHDPE** | LOD> | LOD> | LOD> | LOD> | LOD> | LOD> |
| **7,8-DiHDPE** | LOD> | LOD> | LOD> | LOD> | LOD> | LOD> |
| **17-HDoHE** | LOD> | LOD> | LOD> | LOD> | LOD> | LOD> |
| **22-HDHA** | LOD> | LOD> | LOD> | LOD> | LOD> | LOD> |
| **20-HDHA** | LOD> | LOD> | LOD> | LOD> | LOD> | LOD> |
| **12(13)-EpODE** | LOD> | LOD> | LOD> | LOD> | LOD> | LOD> |
| **12,13-DiHODE** | LOD> | LOD> | LOD> | LOD> | LOD> | LOD> |
| **9(10)-EpODE** | LOD> | LOD> | LOD> | LOD> | LOD> | LOD> |
| **9,10-DiHODE** | LOD> | LOD> | LOD> | LOD> | LOD> | LOD> |
| **13-HOTrE** | LOD> | LOD> | LOD> | LOD> | LOD> | LOD> |
| **9-HOTrE** | LOD> | LOD> | LOD> | LOD> | LOD> | LOD> |
| **8(9)-EpETrE** | LOD> | LOD> | LOD> | LOD> | LOD> | LOD> |
| **8,9-DiHETrE** | LOD> | LOD> | LOD> | LOD> | LOD> | LOD> |
| ***8,15-DiHETE*** | LOD> | LOD> | LOD> | LOD> | LOD> | LOD> |
| **19-HETE** | 169.62 | 155.30 | 267.61 | 150.67 | 117.67 | 135.00 |
| **15-HETE** | 144.42 | 56.97 | 119.70 | 27.33 | 28.83 | 22.10 |
| **12-HETE** | 29.58 | 23.48 | 51.49 | 7.56 | 7.00 | 9.84 |
| **5-HETE** | 23.08 | 15.15 | 20.90 | 0.08 | 0.08 | 0.08 |

| | | | | | | |
|---|---|---|---|---|---|---|
| **15(S)-HETrE** | 105.19 | 67.27 | 109.55 | 44.22 | 46.33 | 42.58 |
| **20-HEPE** | 195.77 | 101.52 | 179.85 | 142.22 | 85.67 | 104.52 |
| **12-HEPE** | 237.31 | 220.61 | 158.66 | 129.11 | 100.67 | 94.52 |

**Solvent optimization for homogenization of worm sample**

*C. elegans* samples were collected, mixed, and aliquoted in 6 equal samples (about 10 mg of worm in each sample). PBS and isopropanol were used separately for solvent optimization of the homogenization step. For each reaction, 100 µl of each solvent was added to equally prepared worm samples (n=3 for each solvent) followed by 10 µL of Type I internal standard (0.4A, concentrations are in SI Table 1) and 10 µl of the antioxidant cocktail including triphenylphosphine (TPP) (0.2 mg/ml), butylated hydroxytoluene (BHT) (0.2 mg/ml), and ethylenediaminetetraacetic (EDTA) (1mg/ml). Two to three stainless steel beads (0.9-2 mm) were added to samples after 10 seconds of mixing by vortex. They were flash frozen in liquid nitrogen and immediately homogenized using an OMNI BEAD RUPTOR homogenizer at a speed of 5.65 Hz for 30 seconds at 4˚C for 3 times. After homogenization, 900 µL of PBS was added to all 6 samples. The samples were centrifuged at 10000g for 5 minutes. After that, the supernatant was collected and was ready for SPE and analysis.

**Solid phase extraction**

Waters Oasis-HLB cartridges (Part No. WAT094226, Lot No. 176A30323A) were used for sample preparation and clean-up. The cartridge was prepared for extraction by washing with ethyl acetate (2mL), methanol (2× 2 mL), and water/methanol (95:5 v/v) with acetic acid 0.1% (2× 2mL). The supernatant of homogenized worm samples (around 10mg), which was spiked before homogenization with 10 µL a 0.4A Type I internal standards solution **(A is defined in SI Table 1)** and 10 µL of antioxidant cocktail as mentioned in solvent optimization, were loaded onto the HLB cartridges. After loading samples, cartridges were washed with water/methanol (95:5 v/v) with acetic acid 0.1% (1.5 mL). The HLB cartridges were dried under a low vacuum for around 20 minutes to remove the residual amount of water and other solvent residues from the cartridge. The HLB cartridges were then eluted with methanol (0.5 mL) followed by ethyl acetate (1 mL) into 2 ml Eppendorf tubes containing glycerol/ MeOH ( 30% v/v, 6 µL) as a trap solution. The eluents were concentrated by the speed vacuum concentrator. The residues were reconstituted in ethanol/water (75% v/v, 100 µL) containing internal standard, CUDA, (10 nM). The samples were then vortex for 5 minutes followed by centrifugal filtration (0.45 µM Millipore) to remove any small particles. The filtrates were transferred to amber vials (1.5 mL capacity) with salinized insert (200 µL capacity), purged with argon gas, caped, and stored at −80°C until UPLC/MS/MS analysis[2].

**LC-MS/MS analysis**

The LC conditions were optimized to separate all eicosanoids of interest with the desired peak shape and signal intensity using an XBridge BEH C18 2.1x150mm HPLC column, ser#01723829118314. Mobile phase A was comprised 0.1% acetic acid in water. Mobile phase B consisted of acetonitrile: methanol (84:16) with 0.1% glacial acetic acid. The gradient elution was performed at a flow rate of 250 µL/min. Chromatography was optimized to separate all analytes in 20 min. The gradient is given in **(SI Table 3 )** less hydrophobic eicosanoids, including PGs, TXS, and LTs were eluted earlier whereas more hydrophobic eicosanoids, including HETEs, HEPEs, AA, and DHA, EPA were eluted later. The autosampler, Waters ACQUITY FTN, was kept at 10°C. The column was connected to a TQXS tandem mass spectrometer by Waters Co. (Milford, MA), equipped with a Waters Acquity SDS pump and Waters Acquity CM detector. Electrospray was operated as an ionization source for negative MRM mode and an interface between HPLC and tandem mass. To achieve the best selectivity and sensitivity, each analyte standard was infused into the mass spectrometer and MRM transitions and source parameters were optimized for that analyte. Analytes were distributed among various periods, which are called functions. In each function, one Type I internal standard was located based on the retention time. The distribution of compounds in different functions increases in dwell time, lowering the limits of detection and increasing the sensitivity. In the whole optimization process, the most abundant transitions were kept for each standard. The more specific or sensitive transitions were selected to distinguish between the close isomers and to gain better detection limits. The optimized mass spectrometric parameters are given in **(Table 1).** All these instrumental optimizations help us to comfort two main issues with oxylipin analysis, low endogenous concentration in biological samples and the similarity in the structure.

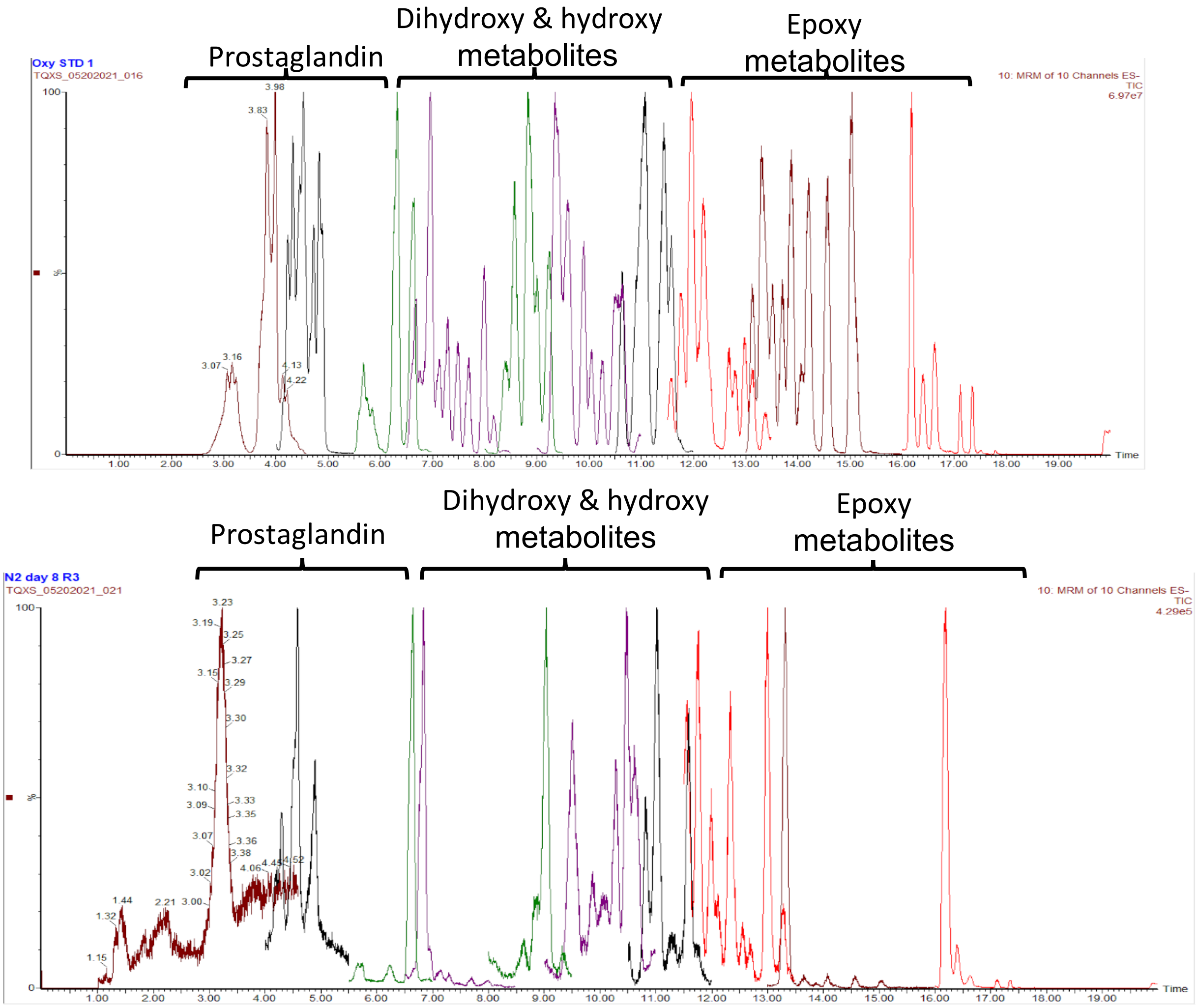

*Figure1. The XIC of PUFAs metabolites from the calibration curve (top) and worm samples (bottom). Multiple reaction monitoring causes a dramatic increase in sensitivity and selectivity in HPLC-MS/MS which allows us to monitor these low-abundance very potent metabolites over the lifespan of C. elegans with minimum samples and cost.*

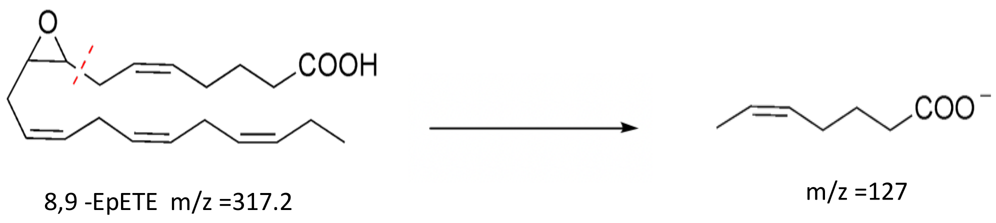

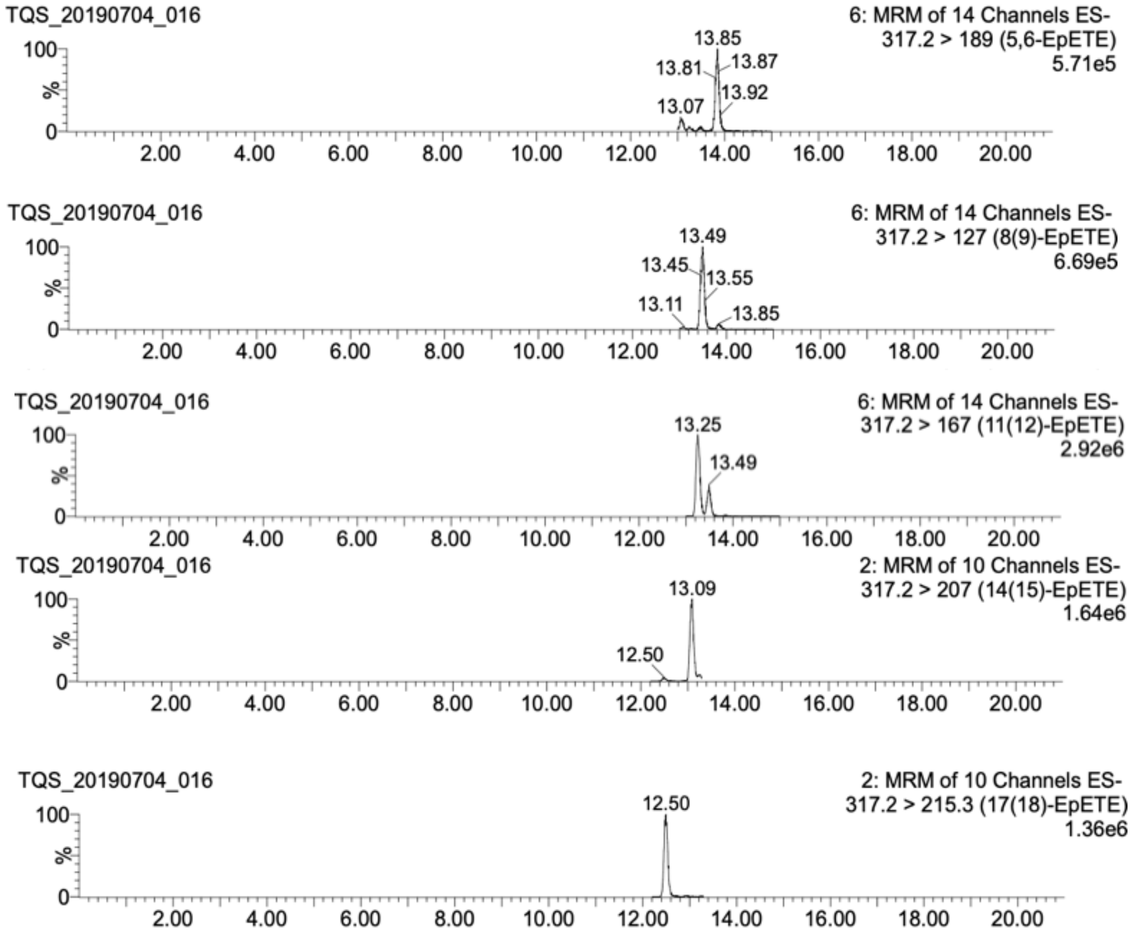

*Figure2. The MRM chromatogram of different isomers of EpETE, demonstrates the separation of five regioisomers of PUFA Metabolite EpETE. This figure illustrates the successful separation of five regioisomers of PUFA metabolite EpETE using a gradient in in the mobile phase and optimized selection of mass transitions in LC-MS/MS analysis. Despite having similar retention times, the distinct mass transitions enable the differentiation of these regioisomers.*

#### Figures of merit

##### LOQ and linearity range

In one day, ten batches of standard mixtures and three further dilutions were injected in four analytical runs back-to-back, to determine the LOQ and linearity range. The reason for adding three more dilutions in the calibration curve was the measurement of the limit of quantification, a signal-to-noise ratio (S/N) >10. The number of spots decreased to eight after the limit of detection was stabilized.

All calibration spots with S/N <10, and deviation >20% were excluded from the calibration curve. The standard concentrations were back-calculated from constructed calibration curves for each analyte, and spots with deviations more than 20% from the nominal concentrations were also excluded. A linear plot of the relation between the concentration and peak area ratio was fitted using 1/x weighting factor linear regression. The correlation coefficient was higher than 0.996 for all oxylipins in the calibration curve (n=3), showing acceptable linearity of the assay in the selected calibration range. The analytes and IS peak retention time were stable between different analytical runs, a 0.2-minute- window was considered as an acceptable difference in retention times.

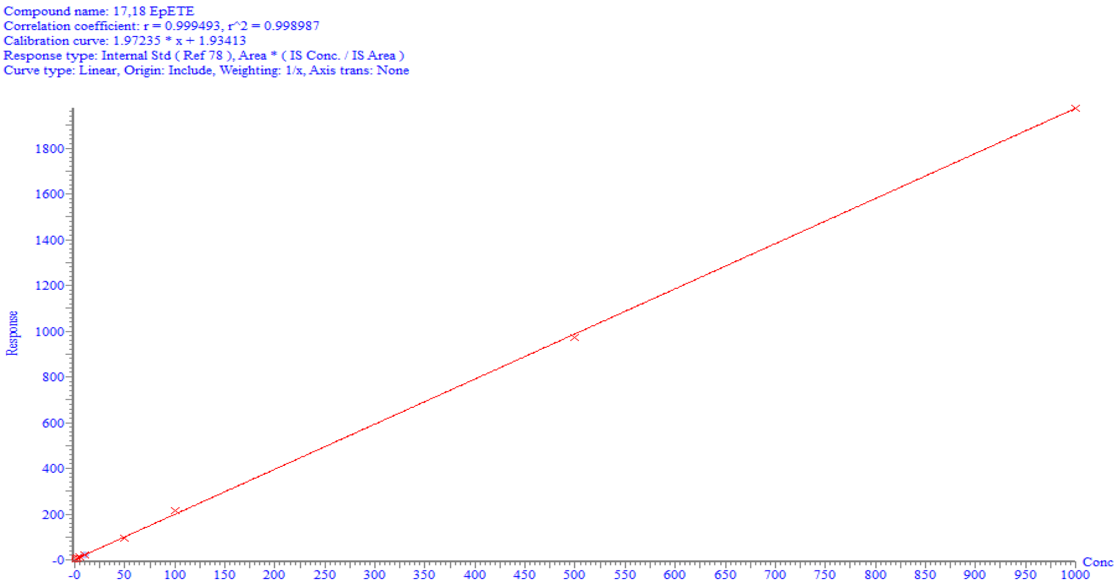

*Figure3. Represents calibration curve of 17,18-EpETE, with r2= 0.998.*

#### Accuracy and precision

Quality control (QC) samples of each analyte were used to calculate the accuracy and precision of the method. These QC samples consisted of four different concentrations of oxylipins 0.5A as High QC, 0.04A as Mid QC, 0.012A as Low QC, and 0.0004A as LLQC (concentration A is defined in **SI Table 2**), they were all diluted in PBS. Five replicates of each QC were extracted by SPE separately and samples were analyzed together with a complete set of calibration standards in three analytical runs, in one day to calculate the intra-day accuracy. An analytical run (n=1) of all QC samples on three consecutive days was used to establish the inter-day accuracy, which was determined as the percent difference between different days. The intra-day accuracy was determined as the percent difference between the mean concentration per analytical run and the expected spiked concentration. The coefficient of variation provided the measure of intra- and inter-day precision. Method recovery determines the amount of analyte spiked in the matrix that can be recovered and quantified after SPE. The PBS solutions spiked with different analyte concentrations were extracted by SPE and analyzed by LC-MS/MS to determine recovery for each analyte. As mentioned in the calibration curve section, a separate calibration curve with different concentrations of Type I internal standard and constant concentration of Type II (0.04A) was used for this calculation.

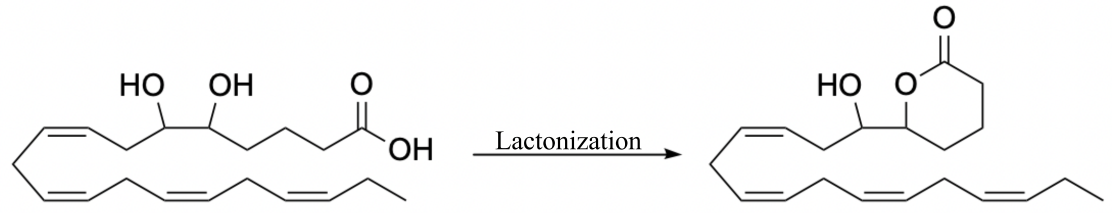

*SI Figure 4. Lactonization of 5,6-DiHETE*

**The matrix effect**

The matrix effect can be determined using the following equation:

$$(ME) = A-B / A*100$$

Where the concentration of Type I IS directly from the calibration curve (EtOH 75%) is denoted by A. Worms sample has undergone sample preparation and spiked with the Type I IS at the same concentration as the calibration curve, which is represented by B. The presence of a matrix effect can be determined by comparing these values. A matrix effect of zero percent indicates the absence of any matrix effect. Conversely, A positive value indicates enhancement, while a negative value implies suppression or interference caused by the matrix.

#### C. *elegans* experiments

##### Epoxide hydrolase inhibitor supplementation

To supplement with the epoxide hydrolase inhibitor, AUDA, we prepared a 20 mM stock solution of AUDA in ethanol. This stock solution was then added into the autoclaved NGM agar solution at a temperature between 55-65°C, resulting in a final concentration of 100 μM. The inhibitor added solution was poured into the petri dishes to make the final treatment plates. The plates were left at room temperature for one day and subsequently inoculated with 250-400 μL of E. coli OP50 (2.8 × 10^8 cells/mL).

##### Age-synchronized worm

Specific numbers of healthy and well-fed Day 1 adult worms were transferred to fresh nematode growth media (NGM) plates (50-100 worms/plate) with OP50 in order to provide the age-synchronized population. The adult worms laid eggs for about 6-10 hours. The eggs were hatched under isolation. About 36-48 hours later, plates were washed off with s-basal solution and transferred to a 40 mm cell strainer placed on top of a 50 mL centrifuge tube. The filtration was used to separate large-sized L4 larvae (stick to the filter), from larva, bacteria carryover, or possible contamination (passed through the filter). L4 larvae were then washed with 75-100 ml of s-basal, transferred to a 1.7 ml centrifuge tube, and centrifuged at 325 x g on a table-top centrifuge for 30 s. The s-basal solution was removed by aspiration, leaving behind a pellet of L4. Finally, L4 worms were resuspended in s-basal solution and transferred to the supplemented or control plates seeded with OP50, recovery of filtration (**Figure 5. SI**).[3]

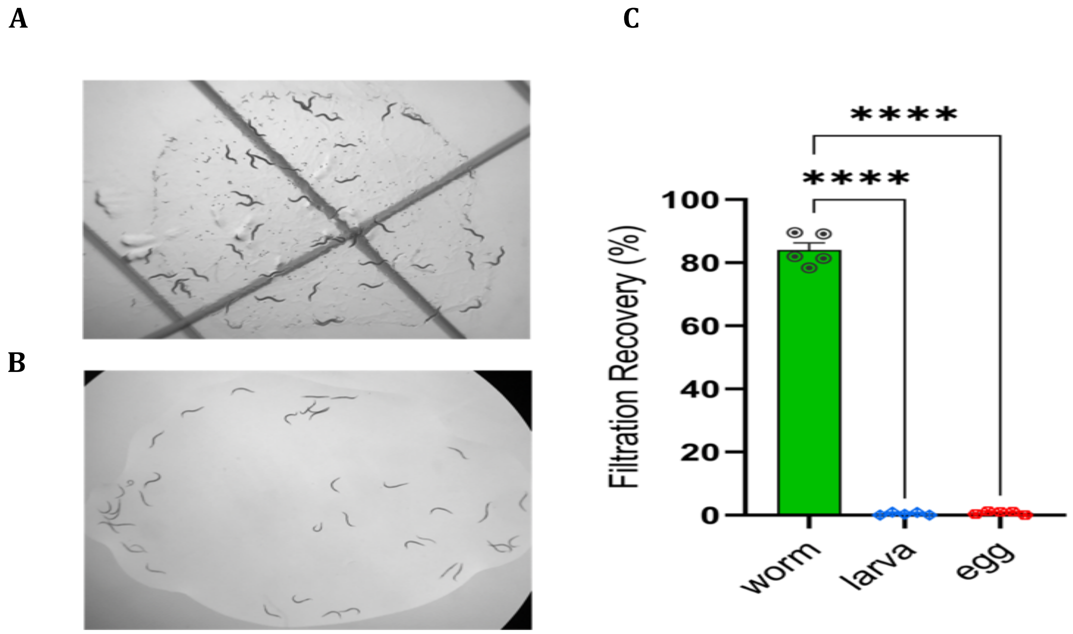

*Figure 5. Filtration step to provide age synchronized worms, A) before filtration, B) after filtration C)filtration recovery % is mean ± SEM (n=5). Statistical differences between worm, larva and egg were evaluated by multiple Student t test, p value >0.0001*
